# *Arabidopsis PIEZO* integrates magnetic field and blue light signaling to regulate root growth

**DOI:** 10.1101/2025.02.11.637623

**Authors:** Ziai Peng, Wenjing Yang, Man Dong, Hanrui Bai, Yan Lei, Ninghui Pan, Yong Xie, Liwei Guo, Changning Liu, Yunlong Du

**Affiliations:** College of Plant Protection, Yunnan Agricultural University, Kunming 650201, China; CAS Key Laboratory of Tropical Plant Resources and Sustainable Use, Yunnan Key Laboratory of Crop Wild Relatives Omics, State Key Laboratory of Plant Diversity and Specialty Crops, Xishuangbanna Tropical Botanical Garden, Chinese Academy of Sciences, Kunming 650223, China; State Key Laboratory for Conservation and Utilization of Bio-Resources in Yunnan, Yunnan Agricultural University, Kunming 650201, China; Key Laboratory of Agro-Biodiversity and Pest Management of Education Ministry of China, Yunnan Agricultural University, Kunming 650201, China; College of Life Sciences, Division of Life Sciences and Medicine, University of Science and Technology of China, Hefei 230026, China

## Abstract

The mechanosensitive ion channel PIEZO is known to play a role in root growth. However, whether the *PIEZO* gene responds to magnetic fields and the mechanisms underlying its regulation of root growth remain unclear. Here, we demonstrate that *Arabidopsis PIEZO* regulates root growth in response to both MF and blue light. Mutation of *PIEZO* led to a significantly shorter roots under MF exposure and blue light illumination. We further identified that *PIEZO* expression in root tips was up-regulated by a blue light-induced signal, which is transmitted from leaves to roots in the presence of a MF. *PIEZO* modulated calcium ion efflux and disturbed auxin transport, specifically through interactions with PIN-FORMED (PIN) 3, 6 and 7 under combined MF and blue light conditions. Notably, the blue light receptors CRYPTOCHROME 1 (CRY1) and CRY2 were essential for both MF perception and the regulation of root growth. Transcriptome analysis of the *piezo-cl* mutant under MF and blue light revealed that the *PIEZO* integrates multiple signaling pathways, including those involved in gibberellin 4 (GA4), ethylene, calcium ion-related genes, mechanosensors, and microRNAs. Specifically, *miR5648-5p* expression conferred MF sensitivity and provided a mechanism for the negative regulation of *PIEZO* under these conditions. Our findings elucidate a multifactorial mechanism by which *PIEZO* coordinates root growth responses to MF and blue light, integrating phytohormone signaling, mechanosensation, calcium ion dynamics, and light perception. This study highlights *PIEZO* as a central node in a complex network that converges diverse environmental cues to regulate root growth.

**One-Sentence Summary:** PIEZO integrates magnetic fields and blue light signaling to regulate root growth in *Arabidopsis* through coordinated phytohormone, calcium, and mechanosensory pathways.

## Introduction

Magnetic fields (MFs) are a fundamental environmental factor essential for organismal survival and development. In plants, MFs play a critical role in growth regulation (*1–4*), including root growth (*5–6*). These effects are mediated through interactions with cryptochromes (CRYs), a class of blue light-sensitive flavoproteins, the auxin signaling pathway (*7–9*), and calcium ion (Ca^2+^) homeostasis (*10–11*). For instance, *Arabidopsis* seedlings grown on the International Space Station, where the galactic MF is 0.1-1 nT (*11*), exhibit altered expression of calcium-related genes compared to Earth-grown seedlings (*12*). Additionally, MF effects on plant growth are linked to the light responses, particularly blue light, which enhances Ca^2+^ influx (*13*) and activates CRY-dependent signaling pathways (*14–16*). Blue light-induced root growth is mediated by CRYs (*17*) and auxin transport that is regulated by PIN-FORMED3 (PIN3) in *Arabidopsis* (*18–19*). Mechanical stress responses in shoot apical meristems (SAMs) involve transient changes in cytoplasmic Ca^2+^ concentrations, which are essential for establishing PIN1 polarity (*20*) and nuclear translocation of the photoreceptor phyB (*21*). Ca²⁺ also plays a central role in root development by modulating auxin signaling (*22–26*). For example, the Ca²⁺ signaling module CALMODULIN IQ-MOTIF CONTAINING PROTEIN (CaM-IQM) interacts with auxin signaling repressors, such as CaM6 binding to IAA19, to regulate root development (*27*). Despite these advances, the interplay among MF, blue light, phytohormones (including auxin), and Ca^2+^ in root growth regulation remains poorly understood.

PIEZO proteins are evolutionarily conserved mechanosensors in land plants (*28*). They form mechanically activated cation channels (*29*) that convert physico-mechanical stimuli into bioelectrical signals (*30*). In *Arabidopsis*, PIEZO localizes to the tonoplast and regulates vacuole morphology (*31*). *AtPIEZO* also functions as a Ca²⁺ channel during mechanical force perception, influencing root growth (*32–33*). The FER-PIF3-PIEZO pathway controls primary root penetration into compacted soil (*34–35*), while the rice *PIEZO* gene contains *cis*-acting elements responsive to light and phytohormones such as methyl jasmonate (MeJA) and gibberellins (GAs) (*36*). These findings suggest that *PIEZO* integrates mechanical, hormonal and light signals to regulate root growth. However, whether *PIEZO* simultaneously responds to MF and blue light to modulate Ca²⁺ homeostasis and auxin transport, thereby regulating root growth, remains unknown. In this study, we demonstrate that *Arabidopsis PIEZO* perceives MF to regulate root growth in a blue light-dependent manner. We reveal that *PIEZO* regulation of root growth integrates multiple signaling pathways involving phytohormones (auxin, GA, and ethylene), Ca²⁺, mechanosensors, and blue light under MF conditions. These finding uncover a new mechanism by which *PIEZO* coordinates root growth through the perception of MF and blue light, highlighting its role as a central regulator of environmental signal integration.

## Results

### *Arabidopsis PIEZO* responds to magnetic field direction in a blue light-dependent manner to regulate root growth

To investigate whether the PIEZO responds to magnetic fields (MFs), we first examined its role in root growth under a 500 mT MF, with the MF direction perpendicular to gravity, in combination with red or blue light (Fig. 1A). When seedlings were grown under red light (Fig. 1B-G), compared with those seedlings not subjected to MF treatment (Fig. 1B, E), the primary root lengths of wild-type (WT) Col-0 (Fig. 1C-D) and *piezo-cl* mutant (Fig. 1F-G) seedlings subjected to a MF were significantly shorter (Fig. 1N) and showed a decreased root growth angle (Fig. 1P-Q). However, when the seedlings were grown under blue light (Fig. 1H-M), compared with the control not subjected to a MF (Fig. 1H, K), the primary root lengths of WT seedlings did not show any obvious change in the presence of a MF (Fig. 1O), but the root lengths of *piezo-cl* mutant seedlings were significantly shorter (Fig. 1O). The root growth angle did not show any obvious change in the WT, but increased in the *piezo-cl* mutant when seedlings were grown in a MF under blue light (Fig. 1R). Similarly, the primary root length and root growth angle of a T-DNA insertion mutant *piezo-T* grown under red or blue light and subjected to MF treatment showed the same phenotype as the *piezo-cl* mutant seedlings (Fig. 1A-R, S1).

**Fig. 1.**
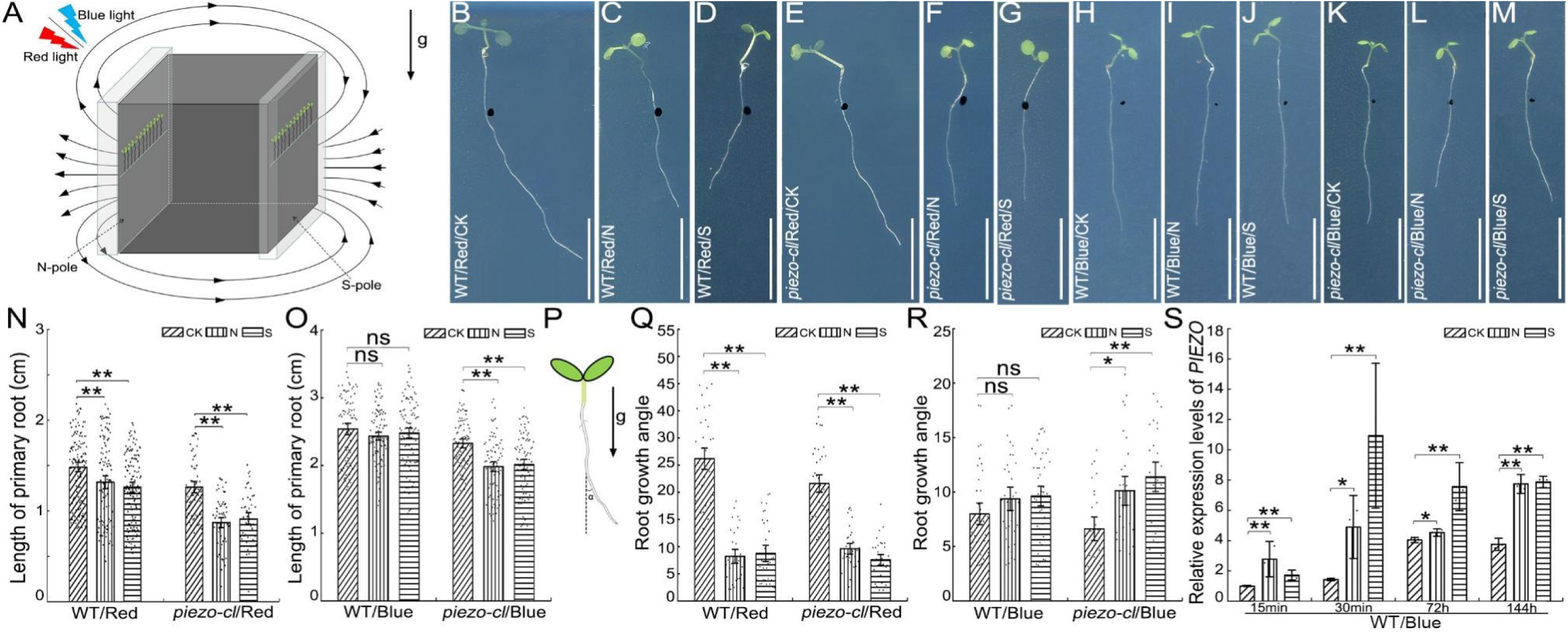
Root phenotypes and *PIEZO* expression in WT and *piezo-cl* mutant seedlings subjected to a MF under red or blue light. Schematic diagram of *Arabidopsis* seedlings subjected to a MF under blue or red light (A). The root phenotypes of WT and *piezo-cl* mutant seedlings treated with 500 mT MF under red light (B-G) and blue light (H-M). Quantification of the primary root length of WT and *piezo-cl* mutant seedlings in a MF under red light (N) (WT: n_CK_ = 120, n_N_ = 105, n_S_ = 127; *piezo-cl*: n_CK_ = 56, n_N_ = 60, n_S_ = 48) or blue light (O) (WT: n_CK_ = 82, n_N_ = 102, n_S_ = 106; *piezo-cl*: n_CK_ = 87, n_N_ = 82, n_S_ = 83). Schematic diagram of root growth angle when seedlings were subjected to a MF showing the root response to gravity (P). Quantification of the root growth angle (α) of WT and *piezo-cl* mutant seedlings under red light (Q) (WT: n_CK_ = 36, n_N_ = 31, n_S_ = 33; *piezo-cl*: n_CK_ = 35, n_N_ = 34, n_S_ = 32) or blue light (R) (WT: n_CK_ = 37, n_N_ = 33, n_S_ = 43; *piezo-cl*: n_CK_ = 26, n_N_ = 29, n_S_ = 26). Expression levels of the *PIEZO* gene in WT seedlings at different times following onset of treatment (15 min, 30 min, 72 hr, and 144 hr) and subjected to a 500 mT MF under blue light (S). The *AtActin2* gene was used as an internal control. Data are means ± SE (Image N-O, Q-R). Arrows labeled “g” indicate the direction of gravity; the black, curved and solid lines with arrow heads represent the direction of the MF lines. The black dotted lines point to the N or S pole of the magnetic block, respectively. WT = wild-type. CK = seedlings not subjected to a MF, N = seedlings treated with the N pole of the magnetic block, S = seedlings treated with the S pole of the magnetic block. Blue = Seedlings treated with blue light. Red = Seedlings treated with red light. Data are means ± SD (Image S), the data presented here represent at least three biological replicates. ns = not significant, * *P* < 0.05, ** *P* < 0.01 (Student’s *t*-test). Scale bar = 1 cm.

When we checked the expression levels of *PIEZO* following different treatment times (15 min, 30 min, 72 hr and 144 hr), we found that the levels of *PIEZO* had increased in the WT seedlings grown under blue light in a 500 mT MF (Fig. 1S). When a transgenic *Arabidopsis* expressing *proPIEZO::GUS* was grown under blue light and with a 500 mT MF, the levels of the GUS gene also increased compared with the control without MF treatment (Fig. S2), confirming that the *PIEZO* perceives MF.

When seedlings overexpressing the *PIEZO* gene were grown under red or blue light in the presence of a MF (Fig. S3A-Y), compared with the control without MF treatment, the WT and *PIEZO*-overexpressing lines #4, #9 and #16 showed reduced root length (Fig. S3Z) and root growth angle (Fig. S3AB) under red light, however, the root length (Fig. S3AA) and root growth angle (Fig. S3AC) did not show significant differences in these lines when grown under blue light.

To assess the effect of MF polarity relative to gravity, WT and *piezo-cl* mutant seedlings were exposed to a 500mT MF, where the direction of the MF was either parallel (Fig. S4A) or anti-parallel (Fig. S4B) to the direction of gravity, under either red or blue light. Compared with the seedlings not subjected to MF treatment, the root lengths of both WT and *piezo-cl* mutant seedlings were significantly reduced when the polarity of the MF was parallel to the direction of gravity under both red (Fig. S4C-F, S) and blue light (Fig. S4G-J, T). However, the root length was not significantly different in the WT and *piezo-cl* mutant seedlings when the polarity of the MF was anti-parallel to the direction of gravity under red light (Fig. S4K-N, U) and blue light (Fig. S4O-R, V). Compared with the root length of WT and *piezo-cl* mutant seedlings grown under blue light was significantly reduced (Fig.S4T) or no difference (Fig.S4V) when the polarity of the MF was parallel or anti-parallel to the direction of gravity, respectively, the root lengths of the WT was unchanged, but root length of *piezo-cl* mutants were significantly shorter when seedlings were grown under blue light and subjected to a MF with polarity perpendicular to the direction of gravity (Fig. 1O). These data indicate that *PIEZO* is sensitive to the polarity of MF under blue light when regulating root growth.

To check the effect of a MF on growth medium, which may then affect seedling root growth, 1/2 MS medium was subjected to a 500 mT MF for 6 days. Compared with seedlings grown in the 1/2 MS medium that was not subjected to a MF (Fig. S5A, D), the root lengths of both the WT (Fig. S5B-C) and *piezo-cl* mutant (Fig. S5E-F) seedlings grown on the 1/2 MS medium previously subjected to a MF did not show significant differences under blue light (Fig. S5G). We next checked whether *piezo* mutant seedling responses to MF regulation of root growth were dependent on the strength of the MF (Fig. S6A). Compared with the seedlings not subjected to a MF (Fig. S6B, E, H, K), the root length of WT (Fig. S6C-D) and *piezo-cl* mutant (Fig. S6F-G) seedlings did not show any significant differences when grown in a 50 mT MF under blue light (Fig. S6N). However, the root lengths were unchanged in the WT (Fig. S6I-J) but significantly reduced in the *piezo-cl* mutant seedlings (Fig. S6L-M) when grown in a 200 mT MF under blue light (Fig. S6O). These data show that the *PIEZO* gene regulates root growth in a MF strength-dependent manner under blue light.

### *PIEZO* perceives a blue light signal transduced from leaves to roots in the presence of a MF

The *PIEZO* promoter contains light-responsive elements (Fig. S7, Table S2). To determine whether *PIEZO* perceives blue light signals transduced from the leaves to the roots, seedlings expressing *ProPIEZO::GUS* were grown with leaves exposed to blue light and roots in darkness under a 500 mT MF (Fig. 2A). Under these conditions, GUS expression was detected in the cotyledon, hypocotyl, vascular tissue and root tip (Fig. 2B-C). Compared with the control seedlings not subjected to a MF (Fig. 2D), the levels of GUS in the root tips (Fig. 2E-F) were higher (Fig. 2G). Similarly, the *PIEZO* expression increased in WT seedlings grown in a MF when compared with the control seedlings were without MF treatment (Fig. 2H). This demonstrates that the *PIEZO* gene is able to perceive the blue light signal transduced from the leaves to roots in the presence of a MF.

**Fig. 2.**
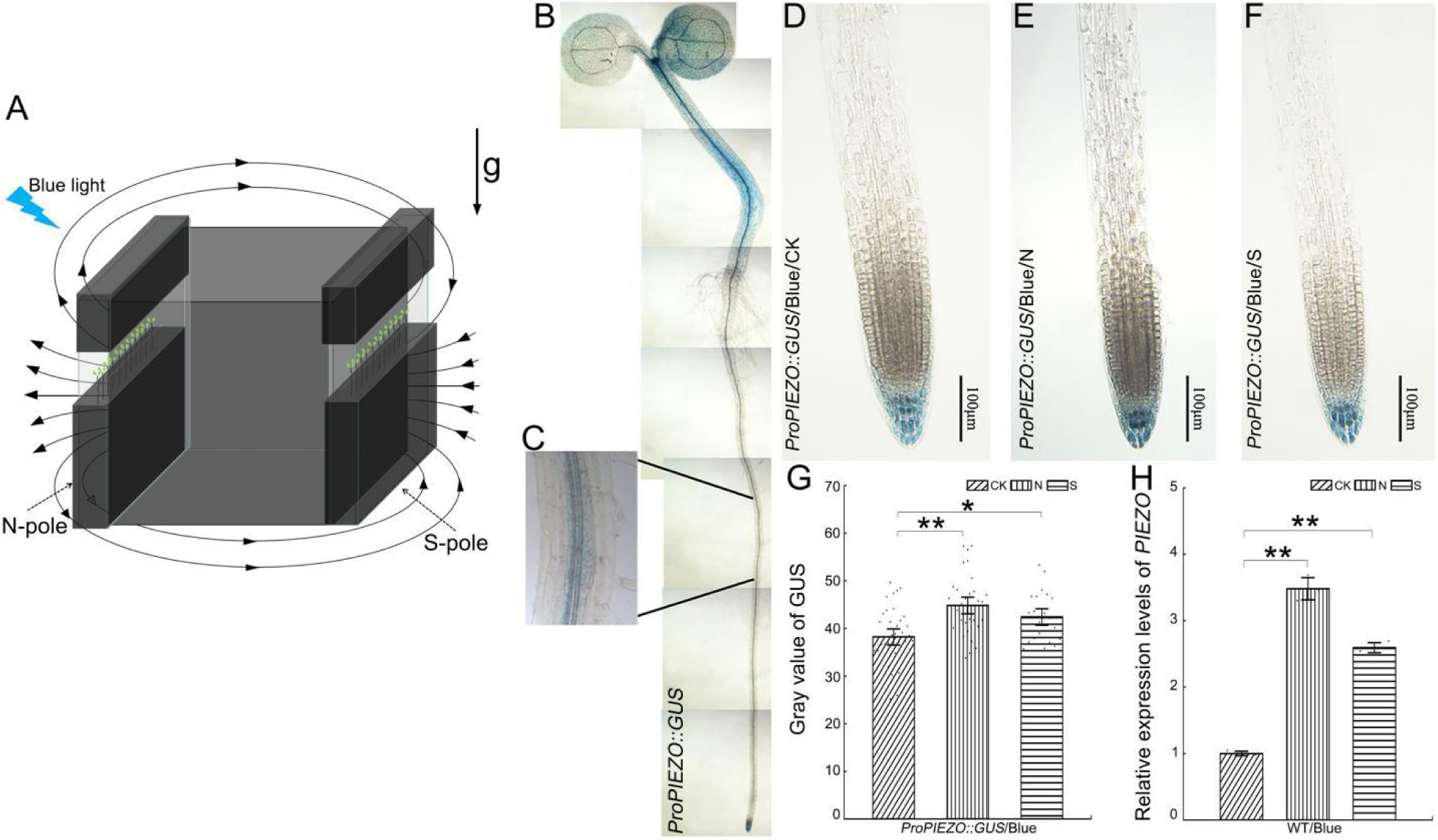
Expression patterns of the *GUS* and *PIEZO* genes in seedlings subjected to a MF under blue light. Schematic diagram of seedlings expressing *ProPIEZO::GUS* subjected to a MF under blue light (A). Expression patterns of GUS in 10-day-old seedlings expressing *ProPIEZO::GUS* (B-C). The GUS expression in the root cap of seedlings expressing *ProPIEZO::GUS* without MF (D) or with 500 mT MF (E-F) under blue light. Quantification of GUS intensity (G) in root tips (n_CK_ = 34, n_N_ = 31, n_S_ = 21). Data are means ± SE (in image G). Expression levels of *PIEZO* gene in the roots of WT seedlings with MF treatment under blue light (H). The *AtActin2* gene was used as an internal control. Arrows labeled “g” indicate the direction of gravity; the black, curved and solid lines with arrow heads represent the direction of the MF lines. The black dotted lines point to the N or S pole of the magnetic block, respectively. WT = wild-type. CK = seedlings not subjected to a MF, N = seedlings treated with the N pole of the magnetic block, S = seedlings treated with the S pole of the magnetic block. Blue = Seedlings treated with blue light. Data are means ± SD (in image H), the data presented here represent at least three biological replicates. ns = not significant, * *P* < 0.05, ** *P* < 0.01 (Student’s *t*-test).

### *PIEZO* regulates Ca^2+^ flux under MF and blue light

The *PIEZO* gene is known to regulate intracellular Ca^2+^ transport in response to mechanical forces (Mousavi et al., 2021). To detect whether *PIEZO* regulation of root growth is related to Ca^2+^ flux, the real-time Ca^2+^ flux in WT and *piezo-cl* mutant seedlings grown under blue light in a 120 mT MF was investigated (Fig. S8). Compared with the control seedlings not subjected to a MF, the Ca^2+^ flux in the roots of WT seedlings was unchanged (Fig. 3A). However, the Ca^2+^ efflux in the roots of *piezo-cl* mutant seedlings (Fig. 3B) grown in a MF under blue light was significantly reduced compared with the control. The above data correlated with root length phenotypes of WT and *piezo-cl* mutant seedlings subjected to a 500 mT MF under blue light (Fig. 1O). To further investigate the effect of MF and blue light on the regulation of Ca^2+^ flux, WT and *piezo-cl* mutant seedlings were grown on 1/2 MS medium (Fig. 3C, D-I) or supplemented with 0.01 μM of the Ca^2+^ inhibitor ethylene glycol bis (2-aminoethyl ether)-N, N, N′, N′-tetraacetic acid (EGTA) (Fig. 3C, J-O) in a 500 mT MF under blue light. The root lengths of WT seedlings grown on 1/2 MS medium with 500 mT MF treatment under blue light did not show difference compared with the seedlings grown in 1/2 MS without a MF treatment (Fig. 3D-F, P).

**Fig. 3.**
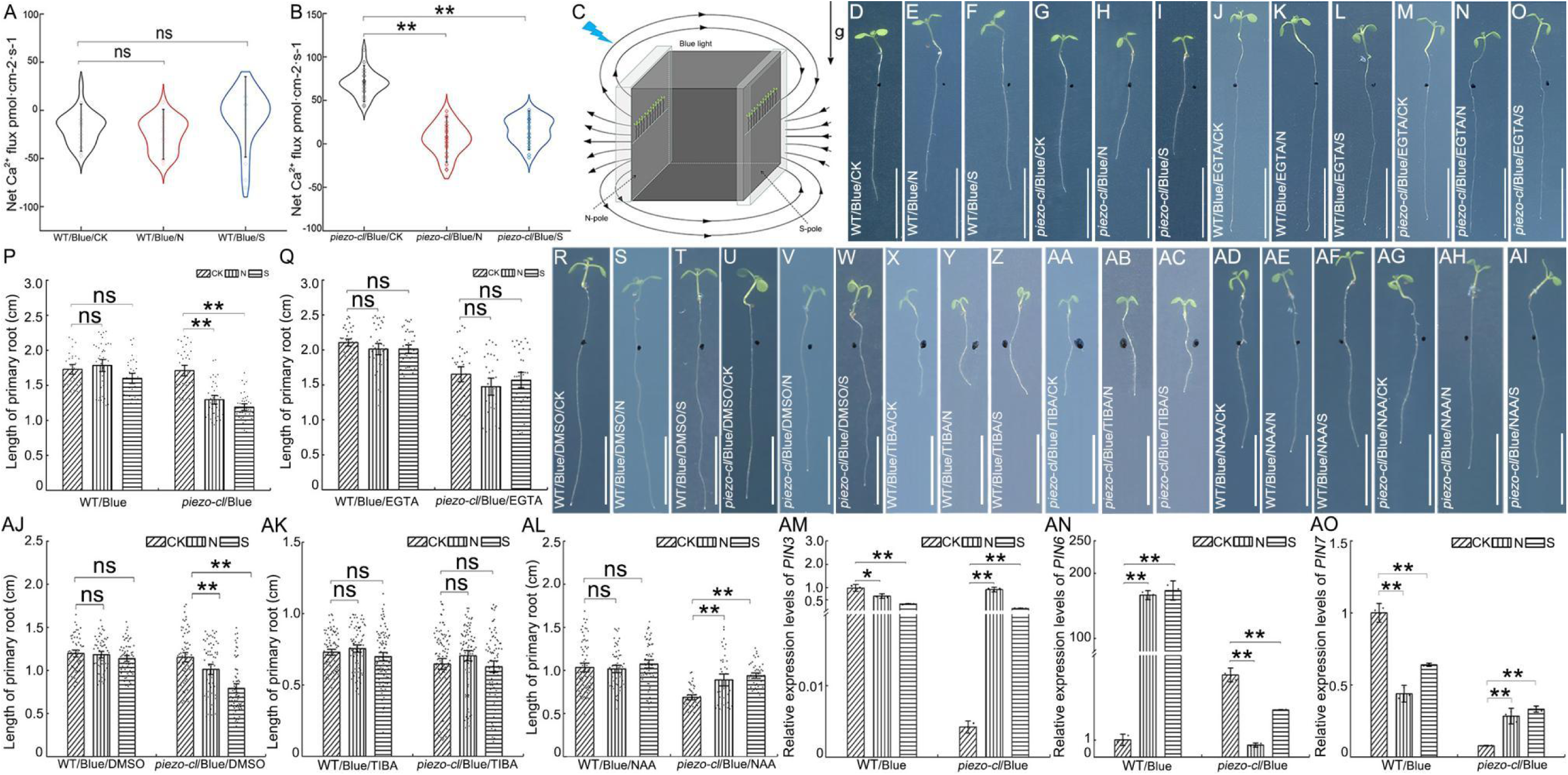
Calcium ion flux in the root tips and root phenotypes of WT and *piezo-cl* mutant seedlings treated with inhibitor of calcium and auxin under MF and blue light treatment. Ca^2+^ flux in the root tips of WT (A) (n_CK_ = 29, n_N_ = 29, n_S_ = 29) and *piezo-cl* mutant (B) seedlings (n_CK_ = 29, n_N_ = 29, n_S_ = 29) subjected to a 500 mT MF under blue light. Data are means ± SD (in image A-B). Schematic diagram of seedlings subjected to MF under blue light (C). The root phenotype of WT (D-F) and *piezo-cl* mutant seedlings (G-I) grown on 1/2 MS medium without MF treatment (D, G) or subjected to a 500 mT MF (E-F, H-I) under blue light (D-I). The root phenotypes of WT (J-L) and *piezo-cl* mutant seedlings (M-O) grown on 1/2 MS medium supplemented with 0.01 μM EGTA but without MF treatment (J, M) or subjected to a 500 mT MF (K-L, N-O) under blue light (J-O). Quantification of the primary root lengths of WT and *piezo-cl* mutant seedlings treated without EGTA as a control (P) (WT: n_CK_ = 28, n_N_ = 37, n_S_ = 32; *piezo-cl*: n_CK_ = 36, n_N_ = 37, n_S_ = 38) or with 0.01 μM EGTA (Q) (WT: n_CK_ = 39, n_N_ = 38, n_S_ = 40; *piezo-cl*: n_CK_ = 32, n_N_ = 29, n_S_ = 34). The root phenotypes of WT (R-T, X-Z, AD-AF) and *piezo-cl* mutant (U-W, AA-AC, AG-AI) seedlings grown on 1/2 MS supplemented with DMSO as a control (R-W), 5 μM TIBA (X-AC) or 0.01 μM NAA (AD-AI) without MF (R, U, X, AA, AD, AG) or 500 mT MF (S-T, V-W, Y-Z, AB-AC, AE-AF, AH-AI) under blue light (R-AI). Quantification of the primary root length of WT and *piezo-cl* mutant seedlings treated with DMSO (AJ) (WT: n_CK_ = 58, n_N_ = 59, n_S_ = 64; *piezo-cl*/Blue/DMSO: n_CK_ = 67, n_N_ = 65, n_S_ = 69), 5 μM TIBA (AK) (WT: n_CK_ = 86, n_N_ = 83, n_S_ = 94; *piezo-cl*: n_CK_ = 97, n_N_ = 87, n_S_ = 86) and 0.01 μM NAA (AL) (WT: n_CK_ = 72, n_N_ = 63, n_S_ = 63; *piezo-cl*: n_CK_ = 46, n_N_ = 40, n_S_ = 47). The expression levels of *PIN3* (AM), *PIN6* (AN) and *PIN7* (AO) genes in WT and *piezo-cl* mutant seedlings subjected to a 500 mT MF under blue light. The *AtActin2* gene was used as an internal control. Data are means ± SD (in images AM-AO). Arrows labeled “g” indicate the direction of gravity; the black, curved and solid lines with arrow heads represent the direction of the MF lines. The black dotted lines point to the N or S pole of the magnetic block, respectively. WT = wild-type. CK = seedlings not subjected to a MF, N = seedlings treated with the N pole of the magnetic block, S = seedlings treated with the S pole of the magnetic block. Blue = Seedlings treated with blue light. Data are means ± SE (in images P-Q, AJ-AL). ns = not significant, ** *P* < 0.01 (Student’s *t*-test). Scale bar = 1 cm.

However, the root lengths of the *piezo-cl* mutant treated with 500 mT MF under blue light were shorter than the control seedlings that without MF treatment (Fig. 3G-I, P). Compared with the WT (Fig. 3J) and *piezo-cl* mutant (Fig. 3M) seedlings that grown on 1/2 MS medium supplemented with 0.01 μM EGTA but without MF treatment under blue light, the primary root lengths of WT (Fig. 3K-L) and *piezo-cl* mutant (Fig. 3N-O) seedlings grown on 1/2 MS medium supplemented with 0.01 μM EGTA with a MF and blue light treatment did not show any significant difference (Fig. 3Q). These results indicate that the PIEZO modulates Ca²⁺ oscillations to regulate root growth under MF and blue light.

### *PIEZO* modulates phytohormone pathway in root growth regulation

The root growth angles of *piezo-cl* mutant seedlings were altered when seedlings were subjected to a MF under blue light (Fig. 1R). To detect whether auxin, which is involved in root growth, has a role in the MF regulation of root growth in the *piezo-cl* mutant, WT and *piezo-cl* mutant seedlings were treated with the auxin transport inhibitor 2, 3, 5-triiodobenzoic acid (TIBA). The root lengths of WT seedlings grown on 1/2 MS medium complemented with DMSO as a control and subjected to a 500 mT MF under blue light (Fig. 3C) were not different from those in seedlings grown without a MF (Fig. 3R-T, AJ), while root lengths in the *piezo-cl* mutant were shorter (Fig. 3U-W, AJ). However, when seedlings were grown on 1/2 MS medium complemented with 5 μM TIBA in a MF under blue light, compared with the control grown without a MF, the primary root length remained unchanged in both the WT and the *piezo-cl* mutant (Fig. 3X-AC, AK). When the seedlings were grown on medium supplemented with 0.01 μM naphthalene-1-acetic acid (NAA) and subjected to a MF under blue light, compared to the control grown without a MF, the primary root length in the WT was unchanged (Fig. 3AD-AF, AL), while it was increased in the *piezo-cl* mutant (Fig. 3AG-AI, AL).

We then checked the expression levels of the auxin efflux carrier genes PIN-FORMED (PIN) 1-8 when seedlings were grown in a 500 mT MF under blue light (Fig. 3AM-AO, S9). Compared with the WT, the levels of *PIN3* (Fig. 3AM) and *PIN7* (Fig. 3AO) were increased in the mutant, but the expression of *PIN6* (Fig. 3AN) was decreased in the roots of *piezo-cl* mutant seedlings subjected to a MF under blue light. This shows that *PIEZO* is able to regulate the intracellular auxin transport mediated by PIN3, 6 and 7 to control root growth in a MF under blue light.

The function of *PIEZO* is connected to ethylene (*35*). Because a MF close to null decreases GA levels (*37*) and because the *PIEZO* promoter contains GA- and MeJA-responsive elements (Fig. S7), we then detected the effects of GA, ethylene and MeJA on the regulation of root growth in *piezo-cl* mutant seedlings subjected to a 500 mT MF under blue light (Fig. S10A). When seedlings were treated with 100 nM 1-aminocyclopropane-1-carboxylic acid (ACC), which is a precursor of ethylene biosynthesis, compared with control seedlings grown on normal 1/2 MS medium and without MF treatment under blue light, no significant difference was detected in the the primary root lengths of WT seedlings, while root lengths were obviously reduced in the *piezo-cl* mutant seedlings subjected to a MF under blue light (Fig. S10B-G, AF). However, the primary root lengths of WT and *piezo-cl* seedlings did not show significant differences compared with the control grown without a MF when seedlings were treated with ACC and MF together under blue light (Fig. S10H-M, AG). Treatment with 0.1 μM GA4 also produced the same root phenotype as ACC (Fig. S10N-Y, AH-AI). However, the primary root lengths of WT and *piezo-cl* seedlings treated with 0.1 μM MeJA and grown in a MF were similar to those of the control treated with DMSO and grown in a MF under blue light (Fig. S10 N-S, Z-AE, AH, AJ).

### CRY1 and 2 can respond to a MF to regulate root growth

The *PIEZO* gene is able to respond to a MF to regulate root growth in the presence of blue light (Fig. 1A-O). The blue light receptors CRY1 and 2 are also known to respond to a MF (*15, 38*) and affect auxin transport in root growth (*7–8*). To detect whether CRY1 and 2 are involved in MF-mediated root growth under blue light, the *cry_1-104_* and *cry_2-1_* single mutants and *cry1/2* double mutant were subjected to a 500 mT MF under blue light (Fig. 4A-M). Compared with the seedlings not subjected to a MF, the primary root lengths of WT seedlings grown in a MF under blue light were unchanged (Fig. 4B-D, N), but root lengths were significantly reduced compared to the control in the *cry_1-104_* (Fig. 4E-G, N) and *cry_2-1_* mutants (Fig. 4H-J, N). However, the primary root lengths of the *cry1/2* double mutant (Fig. 4K-M, N) did not show obvious change compared to the control. This suggests that the blue light signaling pathway is involved in the MF-mediation of root growth, and that the functions of *CRY1* and *2* are redundant in responding to MF. PIF3 is known to interact with PIEZO to regulate primary root growth (*34–35*). When *pif3* mutant seedlings were grown in a 500 mT MF under blue light, similar to the *piezo-cl* mutant (Fig. 1O), the primary root length of the *pif3* mutant was significantly reduced compared with the control grown without a MF (Fig. S11).

**Fig. 4.**
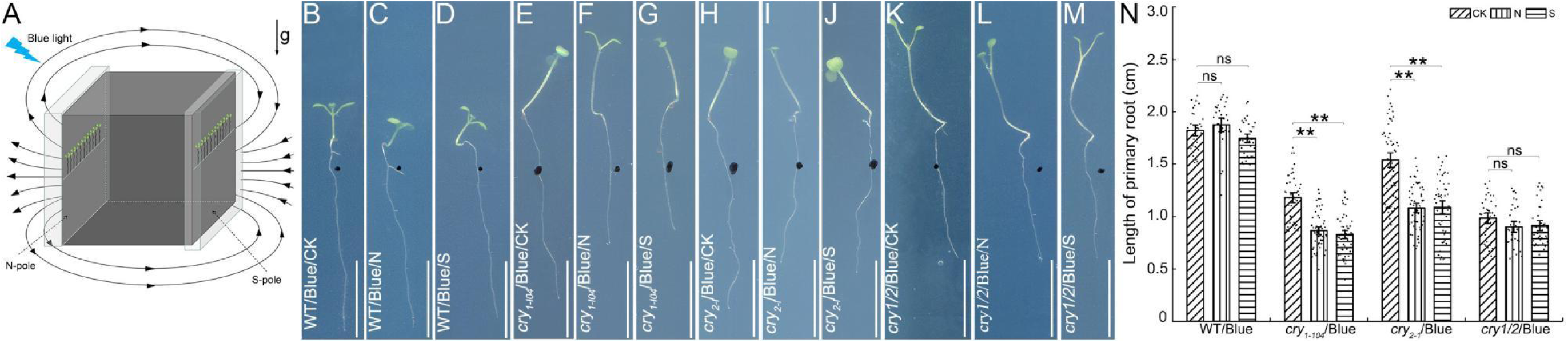
Root phenotypes of *cry1* and *2* mutant line seedlings subjected to a MF under blue light. Schematic diagram of seedlings subjected to a MF under blue light (A). The root phenotypes of WT (B-D), *cry_1-104_* mutant (E-G), *cry_2-1_* mutant (H-J) and *cry1/2* mutant (K-M) seedlings not subjected to a MF (B, E, H, K) or subjected to a 500 mT MF (C-D, F-G, I-J, L-M) under blue light (B-M). Quantification of the primary root length of WT, *cry_1-104,_ cry_2-1_* and *cry1/2* mutant seedlings (N) (WT: n_CK_ = 30, n_N_ = 30, n_S_ = 36; *cry_1-104_*: n_CK_ = 47, n_N_ = 51, n_S_ = 53; *cry_2-1_*: n_CK_ = 55, n_N_ = 56, n_S_ = 54; *cry1/2*: n_CK_ = 36, n_N_ = 34, n_S_ = 37). Arrows labeled “g” indicate the direction of gravity; the black, curved and solid lines with arrow heads represent the direction of the MF lines. The black dotted lines point to the N or S pole of the magnetic block, respectively. WT = wild-type. CK = seedlings not subjected to a MF, N = seedlings treated with the N pole of the magnetic block, S = seedlings treated with the S pole of the magnetic block. Blue = Seedlings treated with blue light. Data are means ± SE, ns = not significant, ** *P* < 0.01 (Student’s *t*-test). Scale bar = 1 cm.

### PIEZO regulates root growth via an integrated gene regulatory network

To gain insight into the mechanism of *PIEZO* regulation of root growth in a MF and under blue light, we analyzed WT and *piezo-cl* seedling root transcriptomes. A total of 1415 differentially expressed genes (DEGs) were found in the root transcriptomes (Table S1, Fig. S12). We found that these DEGs exhibited different expression patterns (Fig. S13), with a total of 1357 genes that specifically responded to blue light were identified in the *piezo-cl* mutant. To investigate the mechanism by which *PIEZO* responds to a MF and blue light, we first constructed an integrated gene regulatory network (iGRN) involving 237 genes by combining transcriptome data from this study, including protein-protein interaction data, miRNA regulation data, and transcription factor regulation data (Fig. S14). Next, we extracted a sub-network closely associated with *PIEZO* (Fig. 5A). Genes from the WT and *piezo-cl* root transcriptomes that had direct or indirect regulatory relationships with *PIEZO*, along with a microRNA m*iR5648-5p*, were identified as being involved in seven distinct biological processes, including auxin signaling, blue light signaling, calcium ion signaling, ethylene signaling, GA signaling, gravitropism, and mechanical pressure (Fig. 5A-B). To further validate the reliability of the dynamic expression changes of gene members in the sub-network, we next checked expression levels of DEGs in the sub-network using real-time PCR. The data showed that expression trends of 8 DEGs (Fig. 5C) were consistent with those of the transcriptome sequencing (Fig. 5B). Additionally, we examined the expression changes of the *miR5648-5p* shown in the sub-network (Fig. 5A), and found that the levels of m*iR5648-5p* were significantly decreased in the roots of WT grown in a 500 mT MF under blue light (Fig. 5D), which showed the opposite expression patterns of the *PIEZO* gene (Fig. 1S). This suggests that this microRNA is able to regulate the expression of *PIEZO* to affect root growth in a MF and under blue light. Based on the iGRN, we propose that *PIEZO* regulation of plant root growth in response to a MF and blue light involves certain phytohormones, including auxin, ethylene, and gibberellin, as well as calcium ion flux, gravity, mechanical pressure and miRNAs.

**Fig. 5.**
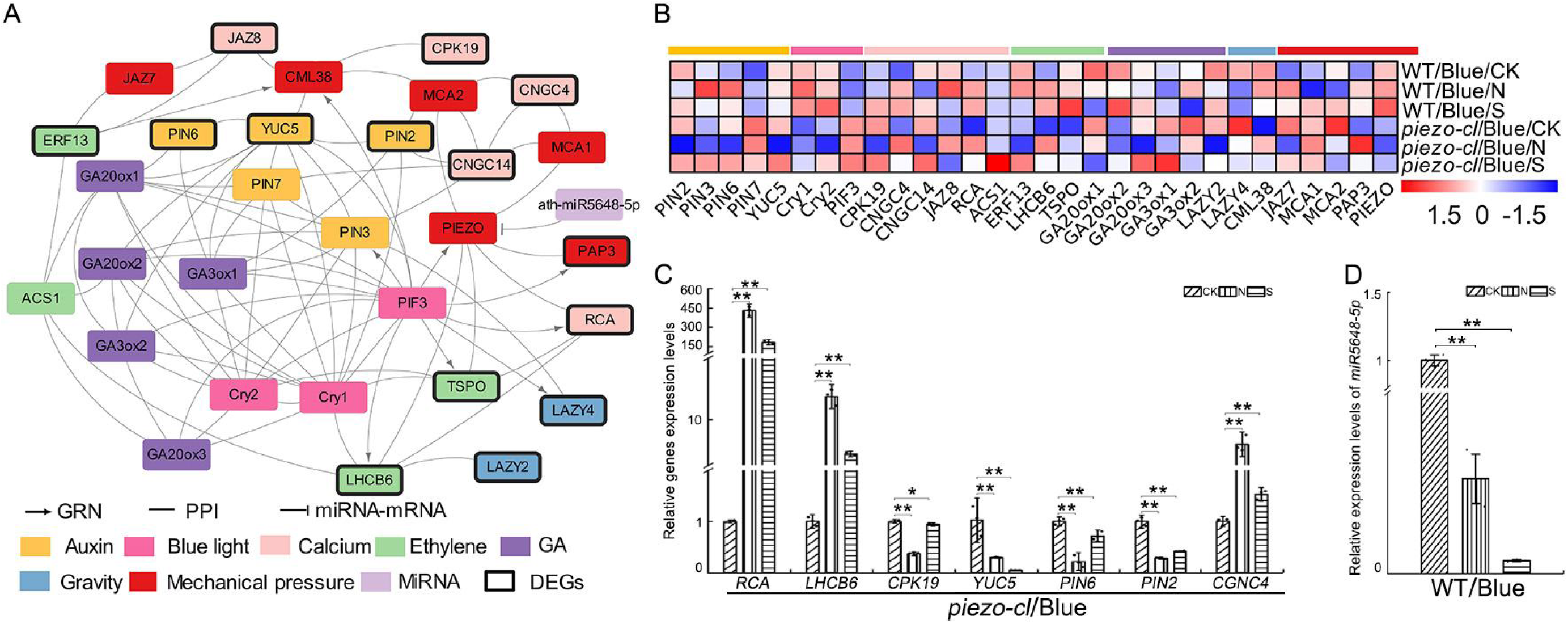
The sub-network of iGRN associated with *PIEZO*. The integrated network of MF-responsive and blue light-responsive genes includes the protein interaction network (PPI), transcription factor regulatory network (GRN) and miRNA-targets regulatory network (miRNA-mRNA) (A). Heatmap showing the normalized FPKM values of unigenes in the sub-network (B). Expression levels of *Arabidopsis* unigenes from the sub-network (C) and *miR5648-5p* (D) in seedlings subjected to a 500 mT MF under blue light. The *AtActin2* gene was used as an internal control. WT = wild-type. CK = seedlings not subjected to a MF, N = seedlings treated with the N pole of the magnetic block, S = seedlings treated with the S pole of the magnetic block. Blue = Seedlings treated with blue light. Data are means ± SD. The data presented here represent at least three biological replicates. * *P* < 0.05, ** *P* < 0.01 (Student’s *t*-test).

## Discussion

The mechanosensitive protein PIEZO plays an important role in plant development. In this study, we demonstrate that the *PIEZO* gene could respond to MF and blue light to regulate root growth in *Arabidopsis*. PIEZO senses the MF polarity relative to gravity, and its expression is induced by an as yet unidentified signal transduced from the leaves to the roots. Under blue light and MF conditions, PIEZO modulates Ca^2+^ flux, phytohormones, including auxin transport, ethylene and gibberellin biosynthesis and signaling pathways, as well as miRNA.

MFs influence plant development, and *PIEZO*-mediated root growth regulation depends on MF polarity—whether perpendicular, parallel, or anti-parallel to gravity (Fig. 1A, S4). Although PIEZO localizes to the vacuole membrane (*31*), the mechanism by which it perceives MF polarity remains unclear. The *PIEZO* gene regulates root growth dependents in blue light (Fig. 1B-O), and the blue light receptors CRY1 and 2 also respond to the MF to modulate root growth (Fig. 4). These findings suggest that PIEZO function within the blue light signaling pathway, as previously proposed (*34–35*), and that either CRY1 or 2 is necessary for the plant to perceive a MF. However, how PIEZO receives and processes blue light signals transduced from the leaves to roots remains unknown.

MF is known to modulate auxin transport to regulate root growth (*7*), and the regulation of root growth by the *PIEZO* gene is associated with Ca^2+^ flux (*33*). Here, we show that PIEZO influenced the Ca*^2+^* efflux and auxin transport mediated by *PIN3*, *PIN6* and *PIN7* (Fig. 3) under MF and blue light conditions. PIEZO localizes to the vascular tissue and the root cap (Fig. 2B-F), overlapping with the expression patterns of PIN3, PIN6, and PIN7 (*39–41*). The iGRN (Fig. 5) includes LAZY2 and LAZY4, which are involved in gravitropic responses (*42*), suggesting that the root cap integrates MF and gravity signals. Ca^2+^ flux, which is closely linked to auxin-mediated root development (*22,25–26*), may be regulated by the PIEZO through mechanosensors such as MCA1 and MCA2, or the calcium-related protein RCA, either directly or indirectly, and may further modulate auxin transport via ethylene-related genes.

The *PIEZO* regulates plant development through interactions with multiple phytohormones (*36*). We found that GA and ethylene are involved in PIEZO-mediated root growth regulation under MF and blue light (Fig. S10). PIEZO did not directly regulate or interact with the GA-related genes, except for ethylene-related genes (Fig. 5A). Although near-null MFs alter the GA-related gene expression (*37*), no differential expression of the GA-related genes was observed in the *piezo-cl* mutant (Fig. 5A). The effect of GA4 on root growth in the *piezo-cl* mutant under MF and blue light (Fig. S10AH-AI) may involve an unidentified pathway or reflect MF-induced changes in GA-related gene expression linked to *PIEZO* mutation. Ethylene may regulate root growth in the *piezo-cl* mutant via the PIF3, as previously reported (*35*). Notably, the miRNA expression was also modulated by MF and could negatively regulate the *PIEZO* expression (Fig. 5A, D), suggesting that certain upstream regulatory elements contribute to PIEZO-mediated root growth under MF and blue light.

PIEZO not only perceives mechanical forces but also responds to MF and blue light. Future studies should focus on elucidating how PIEZO detects MF polarity and its connection to CRY1 and CRY2. Additionally, the mechanism by which miRNAs sense MFs and regulate *PIEZO* expression warrants further investigation. This study reveals a new mechanism by which *PIEZO* integrates MF and blue light signals to regulate root growth, providing new insights into how the Earth’s magnetic field and light environment collectively shape plant development.

## Acknowledgments

We thank Professor Kai He (Lanzhou University) for her generous provision of the *piezo-cl* mutant and the transgenic *Arabidopsis* line expressing *ProPIEZO::GUS*. We also thank Professor Shuhua Yang (China Agricultural University) for the gift of the *cry_1-104_* and *cry_2-1_* mutant lines. Our thanks also go to Professor Haodong Chen (Tsinghua University) for provision of the *cry2/1* double mutant line.

## Author contributions

Y.D. initiated the project, Y.D. and C.L. supervised the project and designed the experiments; Z.P., W.Y. and M.D. performed the majority of the experiments and analyzed and prepared the figures; H.B., Y.L., N.P., Y.X. and L.G. performed additional experiments. Y.D., C.L., Z.P., W.Y., M.D. and H.B. analyzed the data. Y.D. and C.L. wrote and revised the paper with input from all authors.

## Data and materials availability

All data are available in the manuscript or the supplementary materials. The raw data of transcriptomes have been uploaded to the Genome Sequence Archive (GSA) public database (https://ngdc.cncb.ac.cn/) under project number PRJCA027762.

**SUPPLEMENTARY MATERIALS**

**Materials and Methods**

**Figs. S1 to S14**

**Tables S1 to S3**

**References (43 - 60)**

## SUPPLEMENTARY MATERIALS

**Fig. S1.**
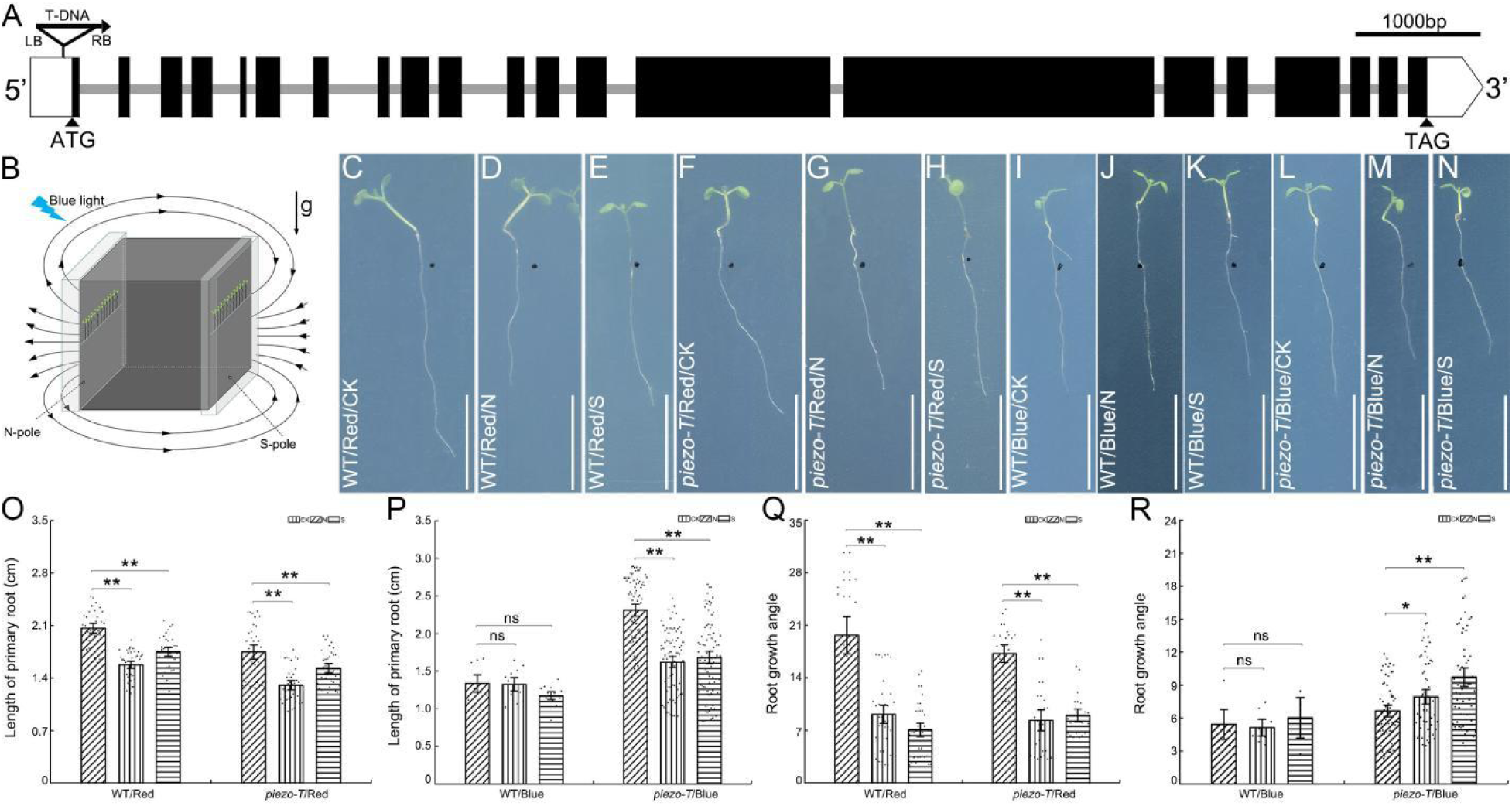
Root phenotypes of *piezo-T* mutant seedlings subjected to a MF under blue light. Schematic diagram of the *PIEZO* gene with a T-DNA insertion (A). Schematic diagram of seedlings subjected to a MF under blue light (B). The root phenotypes of WT (C-E, I-K) and *piezo-T* mutant (F-H, L-N) seedlings without a MF (C, F, I, L) or with a 500 mT MF treatment (D-E, G-H, J-K, M-N) under red light (C-H) or blue light (I-N). Quantification of the primary root length of WT and *piezo-T* mutant seedlings under MF treatment and grown under red light (O) (WT: n_CK_ = 32, n_N_ = 36, n_S_ = 33; *piezo-T*: n_CK_ = 33, n_N_ = 35, n_S_ = 34) or blue light (P) (WT: n_CK_ = 10 n_N_ = 12, n_S_ = 13; *piezo-T*: n_CK_ = 73, n_N_ = 78, n_S_ = 80). Quantification of the root growth angle (α) of WT and *piezo-T* mutant seedlings under MF treatment and grown under red light (Q) (WT: n_CK_ = 24, n_N_ = 34, n_S_ = 36; *piezo-T*: n_CK_ = 24, n_N_ = 25, n_S_ = 22) or blue light (R) (WT: n_CK_ = 6, n_N_ = 8, n_S_ = 4; *piezo-T*: n_CK_ = 60, n_N_ = 56, n_S_ = 58). Black boxes indicate exons, white boxes indicate 5’ UTR and 3’ UTR, gray lines indicate introns, triangle indicates T-DNA insertion sites and the arrow head shows the direction of T-DNA insertion, and the filled triangles indicate the initiation codon ATG and termination codon TAG, respectively. Arrows labeled “g” indicate the direction of gravity; the black, curved and solid lines with arrow heads represent the direction of the MF lines. The black dotted lines point to the N or S pole of the magnetic block, respectively. LB = the left border of T-DNA, RB = the right border of T-DNA. WT = wild-type. CK = seedlings not subjected to a MF, N = seedlings treated with the N pole of the magnetic block, S = seedlings treated with the S pole of the magnetic block. Blue = Seedlings treated with blue light. Data are means ± SE. ns = not significant, * *P* < 0.05, ** *P* < 0.01 (Student’s *t*-test). Scale bar = 1 cm.

**Fig. S2.**
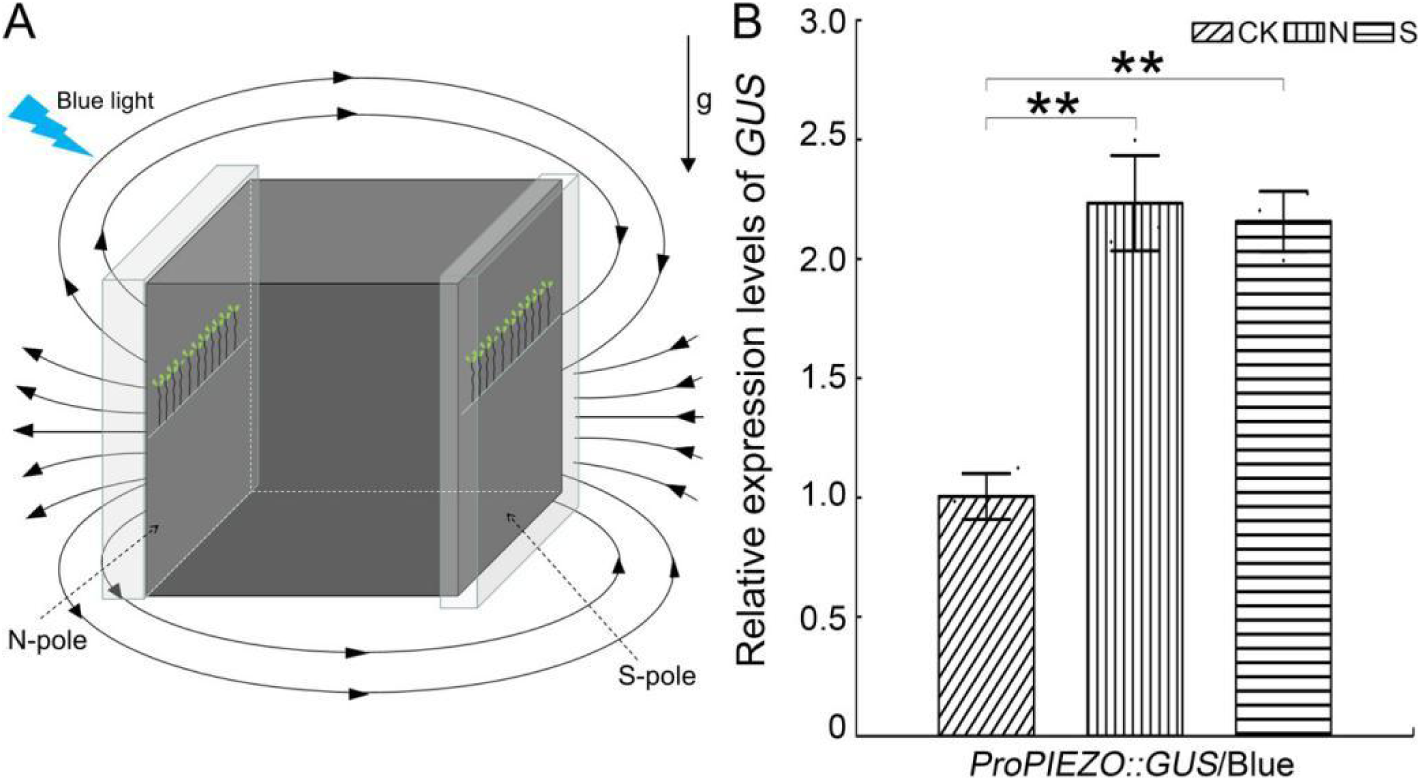
Expression levels of *GUS* gene in *Arabidopsis* seedlings subjected to a MF under blue light. Schematic diagram of seedlings subjected to a MF under blue light (A). (B) Expression levels of the *GUS* gene in the roots of seedlings expressing *ProPIEZO::GUS* in a 500 mT MF under blue light. The *AtActin2* gene was used as an internal control. Arrows labeled “g” indicate the direction of gravity; the black, curved and solid lines with arrow heads represent the direction of the MF lines. The black dotted lines point to the N or S pole of the magnetic block, respectively. WT = wild-type. CK = seedlings not subjected to a MF, N = seedlings treated with the N pole of the magnetic block, S = seedlings treated with the S pole of the magnetic block. Blue = Seedlings treated with blue light. Data are means ± SD. The data presented here represent at least three biological replicates. ** *P* < 0.01 (Student’s *t*-test).

**Fig. S3.**
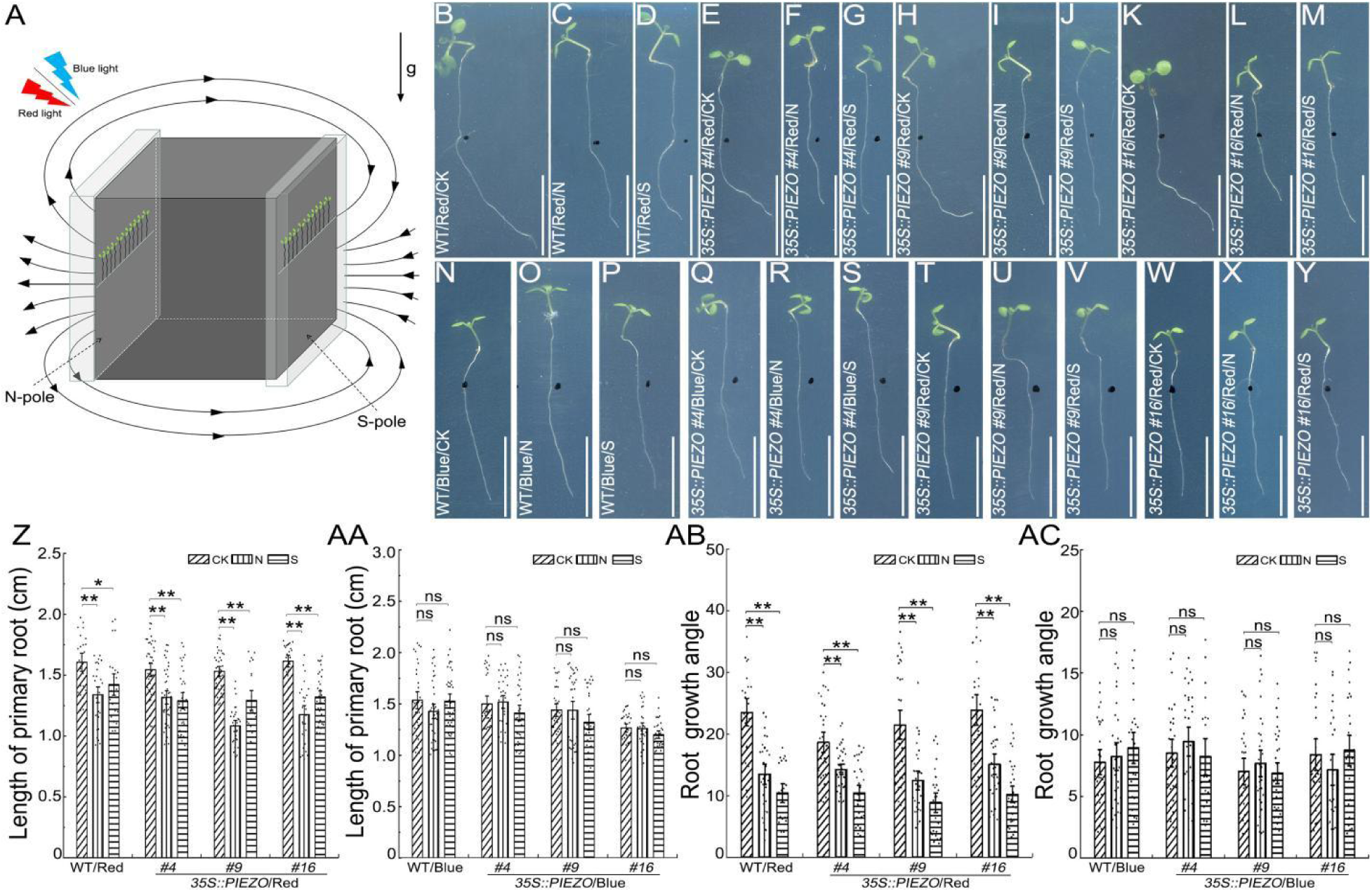
Root phenotypes of seedlings from *35S::PIEZO* lines in a MF under red or blue light. Schematic diagram of seedlings subjected to a MF under blue or red light (A). The root phenotypes of WT (B-D, N-P) and *35S::PIEZO* lines #4 (E-G, Q-S), #9 (H-J, T-V), #16 (K-M, W-Y) not subjected to a MF (B, E, H, K, N, Q, T, W) or in a 500 mT MF (C-D, F-G, I-J, L-M,O-P, R-S, U-V, X-Y) under red light (B-M) or blue light (N-Y). Quantification of the primary root length of WT and *35S::PIEZO* lines #4, #9, #16 subjected to a MF under red light (Z) (WT: n_CK_ = 23, n_N_ = 30, n_S_ = 23; #4: n_CK_ = 41, n_N_ = 42, n_S_ = 33; #9: n_CK_ = 36, n_N_ = 29, n_S_ = 21; #16: n_CK_ = 24, n_N_ = 26, n_S_ = 33) or blue light (AA) (WT: n_CK_ = 33, n_N_ = 39, n_S_ = 41; #4: n_CK_ = 28, n_N_ = 34, n_S_ = 35; #9: n_CK_ = 31, n_N_ = 37, n_S_ = 33; #16: n_CK_ = 31, n_N_ = 29, n_S_ = 32). Quantification of the root growth angle (α) in seedlings of WT and *35S::PIEZO* lines #4, #9, #16 subjected to a MF under red light (AB) (WT: n_CK_ = 22, n_N_ = 27, n_S_ = 21; #4: n_CK_ = 31, n_N_ = 36, n_S_ = 30; #9: n_CK_ = 32, n_N_ = 29, n_S_ = 26; #16: n_CK_ = 23, n_N_ = 31, n_S_ = 34) or blue light (AC) (WT: n_CK_ = 27, n_N_ = 30, n_S_ = 25; #4: n_CK_ = 28, n_N_ = 31, n_S_ = 21; #9: n_CK_ = 29, n_N_ = 32, n_S_ = 33; #16: n_CK_ = 29, n_N_ = 26, n_S_ = 28). Arrows labeled “g” indicate the direction of gravity; the black, curved and solid lines with arrow heads represent the direction of the MF lines. The black dotted lines point to the N or S pole of the magnetic block, respectively. WT = wild-type. CK = seedlings not subjected to a MF, N = seedlings treated with the N pole of the magnetic block, S = seedlings treated with the S pole of the magnetic block. Blue = Seedlings treated with blue light. Red = Seedlings treated with red light. Data are means ± SE. ns = not significant, * *P* < 0.05, ** *P* < 0.01 (Student’s *t*-test).

**Fig. S4.**
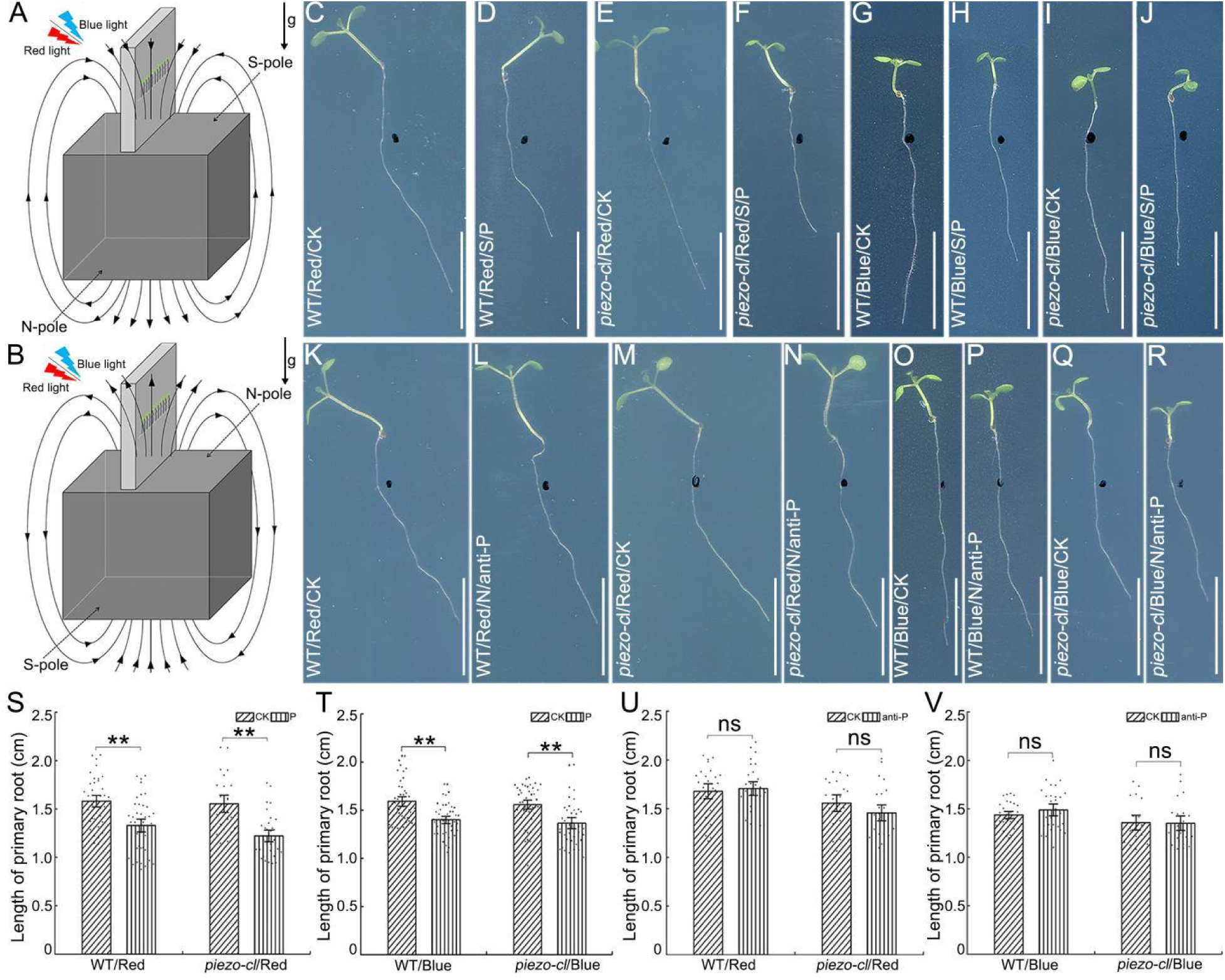
Root phenotypes of *piezo-cl* mutant seedlings subjected to a MF in parallel and anti-parallel with gravity under red or blue light. Schematic diagram of seedlings subjected to a MF in parallel (A) or anti-parallel (B) with the direction of gravity under blue or red light. The root phenotypes of WT (C-D, G-H, K-L, O-P) and *piezo-cl* mutant (E-F, I-J, M-N, Q-R) seedlings subjected to a 500 mT MF parallel to the direction of gravity (C-J) or anti-parallel to the direction of gravity (K-R) under red light (C-F, K-N) or blue light (G-J, O-R). (S) Quantification of the primary root length of WT and *piezo-cl* mutant seedlings subjected to a MF in parallel to the direction of gravity under red light (S) (WT: n_CK_ = 39, n_G_ = 40; *piezo-cl*: n_CK_ = 24, n_G_ = 32) or blue light (T) (WT: n_CK_ = 42, n_G_ = 45; *piezo-cl*: n_CK_ = 49, n_G_ = 38). Quantification of the primary root length of WT and *piezo-cl* mutant seedlings subjected to a MF in anti-parallel to the direction of gravity under red light (U) (WT: n_CK_ = 26, n_Anti-G_ = 27; *piezo-cl*: n_CK_ = 20, n_Anti-G_ = 24) or blue light (WT: n_CK_ = 29, n_Anti-G_ = 29; *piezo-cl*: n_CK_ = 19, n_Anti-G_ = 21). Arrows labeled “g” indicate the direction of gravity; the black, curved and solid lines with arrow heads represent the direction of the MF lines. The black dotted lines point to the N or S pole of the magnetic block, respectively. WT = wild-type. CK = seedlings not subjected to a MF, N = seedlings treated with the N pole of the magnetic block, S = seedlings treated with the S pole of the magnetic block. Blue = Seedlings treated with blue light. Red = Seedlings treated with red light. P = the direction of the MF is parallel to the direction of gravity, anti-P = the direction of MF is anti-parallel to the direction of gravity. Data are means ± SE. ns = not significant, * *P* < 0.05, ** *P* < 0.01 (Student’s *t*-test). Scale bar = 1 cm.

**Fig.S5.**
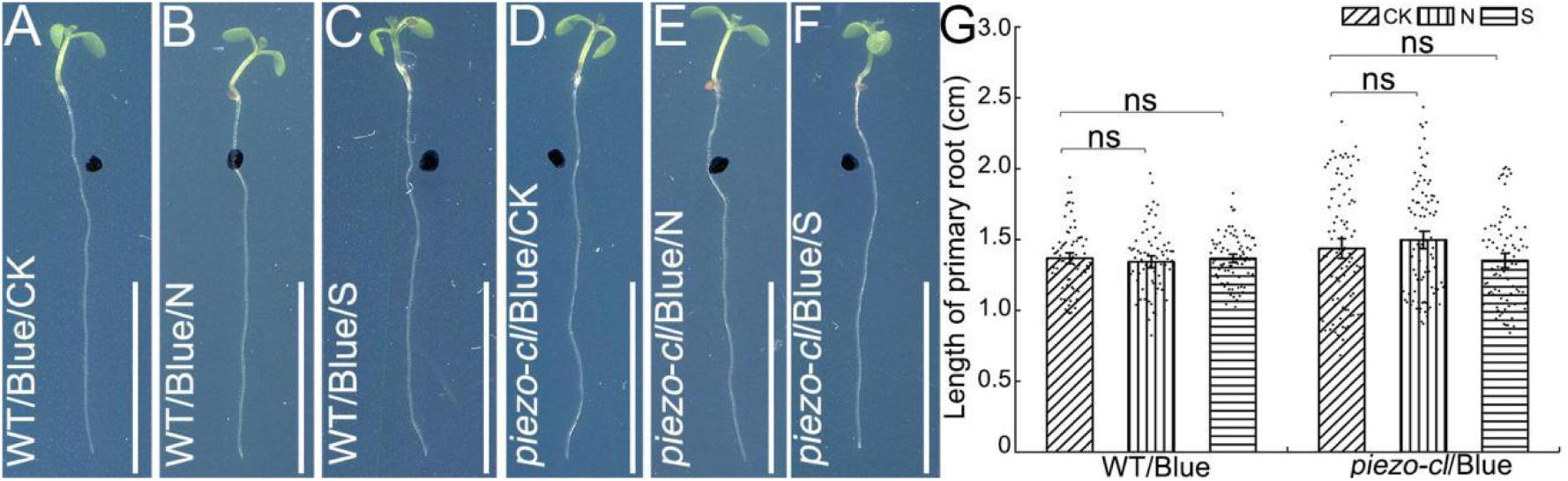
Root phenotypes of *piezo-cl* mutant seedlings grown on MF pre-treated medium under blue light. Root phenotypes of 6-day-old seedlings of WT (A-C) and *piezo-cl* mutant (D-F) seedlings grown on 1/2 MS medium without MF treatment (A, D) or pre-treated with 500 mT MF (B-C, E-F) under blue light (A-F). Quantification of the primary root length of WT and *piezo-cl* mutant (G) seedlings (WT: n_CK_ = 76, n_N_ = 70, n_S_ = 77; *piezo-cl*: n_CK_ = 92, n_N_ = 91, n_S_ = 85). WT = wild-type. CK = 1/2 MS medium without MF treatment, N = 1/2 MS medium was pre-treated with the N pole of the magnetic block, S = 1/2 MS medium was pre-treated with the S pole of the magnetic block. Blue = Seedlings treated with blue light. Data are means ± SE. ns = not significant. Scale bar = 1 cm.

**Fig.S6.**
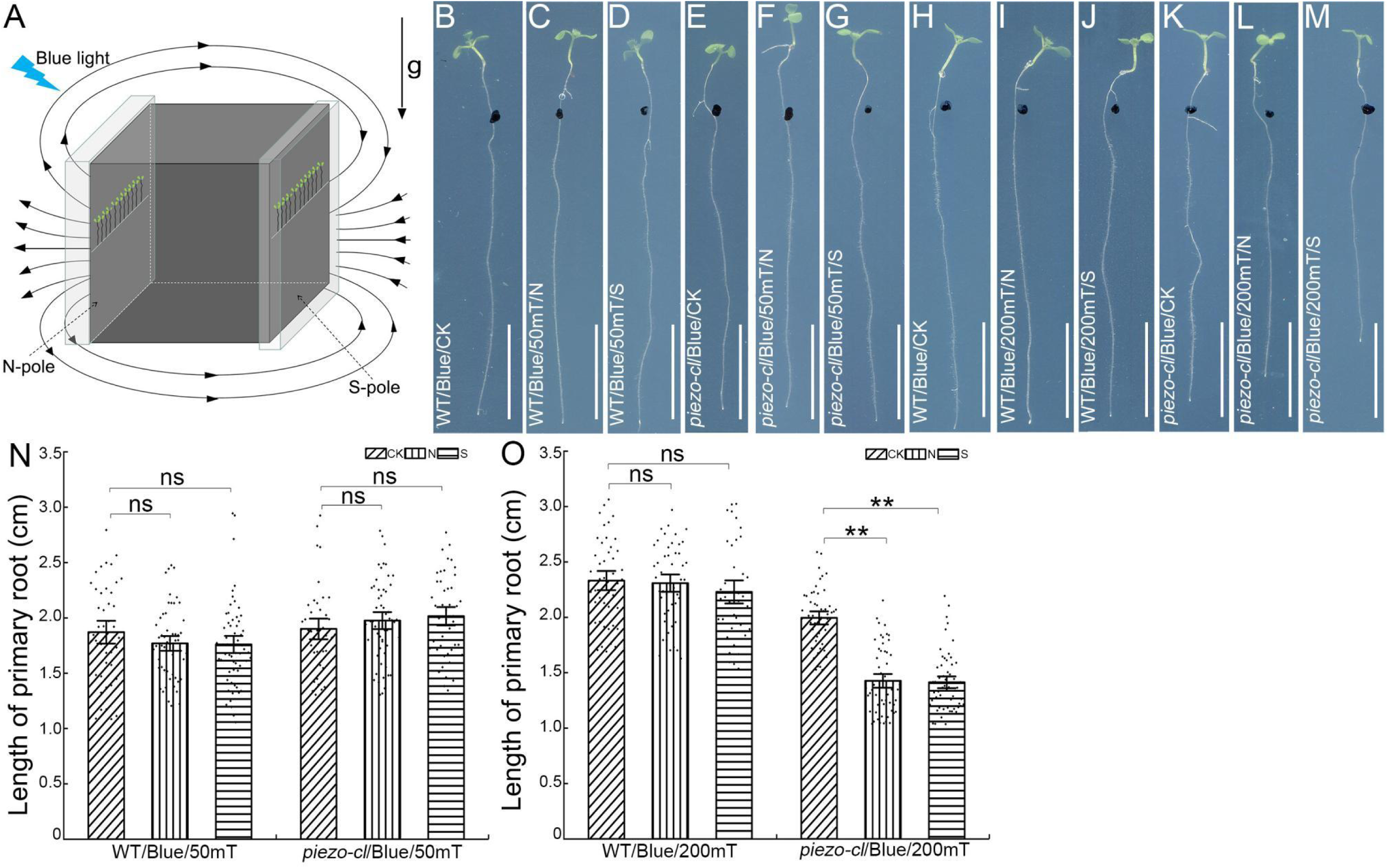
Root phenotypes of *piezo-cl* mutant seedlings subjected to different MF intensities under blue light. Schematic diagram of seedlings subjected to a MF under blue light (A). The root phenotypes of 6-day-old seedlings of WT (B-D, H-J) and *piezo-cl* mutant lines (E-G, K-M) not subjected to a MF (B, E, H, K) or subjected to a 50 mT MF (C-D, F-G) or a 200 mT MF (I-J, L-M) under blue light (B-M). Quantification of the primary root length of WT and *piezo-cl* mutant seedlings subjected to a 50 mT MF (N) (WT: n_CK_ = 43, n_N_ = 54, n_S_ = 58; *piezo-cl*: n_CK_ = 40, n_N_ = 54, n_S_ = 46) or a 200 mT MF (O) (WT: n_CK_ = 44, n_N_ = 48, n_S_ = 39; *piezo-cl*: n_CK_ = 50, n_N_ = 53, n_S_ = 57). Arrows labeled “g” indicate the direction of gravity; the black, curved and solid lines with arrow heads represent the direction of the MF lines. The black dotted lines point to the N or S pole of the magnetic block, respectively. WT = wild-type. CK = seedlings not subjected to a MF, N = seedlings treated with the N pole of the magnetic block, S = seedlings treated with the S pole of the magnetic block. Blue = Seedlings treated with blue light. Data are means ± SE. ns = not significant, ** *P* < 0.01 (Student’s *t*-test). Scale bar = 1 cm.

**Fig.S7.**
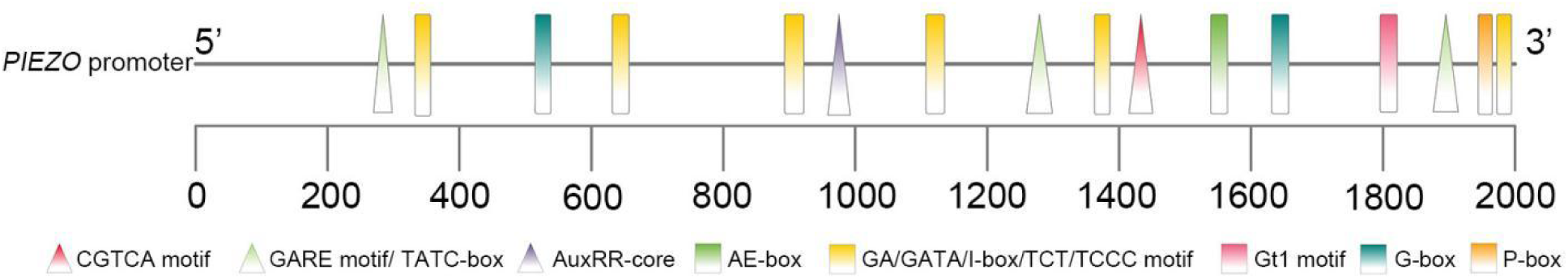
Structure of the *Arabidopsis PIEZO* promoter. The *Arabidopsis PIEZO* promoter contains eight light responsive elements comprising an AE-box (AGAAACTT), GA motif (ATAGATAA), GATA motif (AAGGATAAGG), I-box motif (GGATAAGGTG), TCT motif (TCTTAC), TCCC motif (TCTCCCT), Gt1 motif (GTGTGTGAA), and G-box (TACGTG/CACGAC); an auxin responsive element comprising an AuxRR-core (GGTCCAT); three GA responsive element comprising a P-box (CCTTTTG), GARE motif (TCTGTTG), TATC-box (TATCCCA); and a MeJA responsive element comprising a CGTCA-motif (CGTCA).

**Fig.S8.**
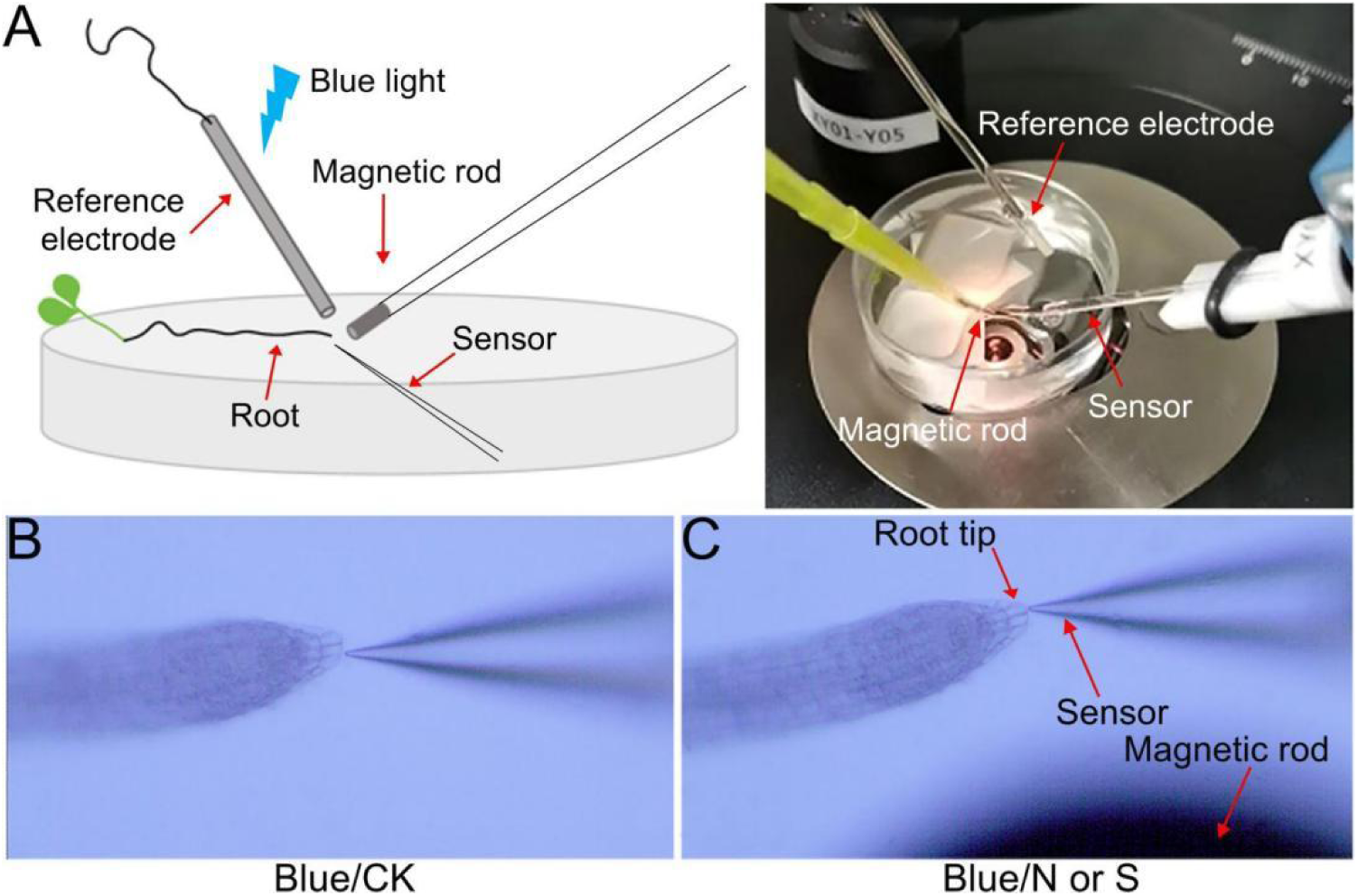
Schematic diagram of calcium ion measurement in the root tips of *piezo-cl* mutant seedlings subjected to a MF under blue light. The roots of seedlings of WT and *piezo-cl* mutants were fixed in 0.1 mM CaCl_2_ solution and treated with the 120 mT magnetic rod under blue light. The Ca^2+^ flux rate in the root tip was determined in real-time using a sensor (A). The Ca^2+^ flux in the root tips of seedlings not subjected to MF (B) or treated with a magnetic rod (C) under blue light and detected using a sensor. The black zone is the image of magnetic rod seen under the microscope. Blue = Roots of seedlings treated with blue light. WT = wild-type. CK = Roots of seedlings without magnetic rod treatment, N = Roots of seedlings treated with the N pole of the magnetic rod, S = Roots of seedlings treated with the S pole of the magnetic rod.

**Fig. S9.**
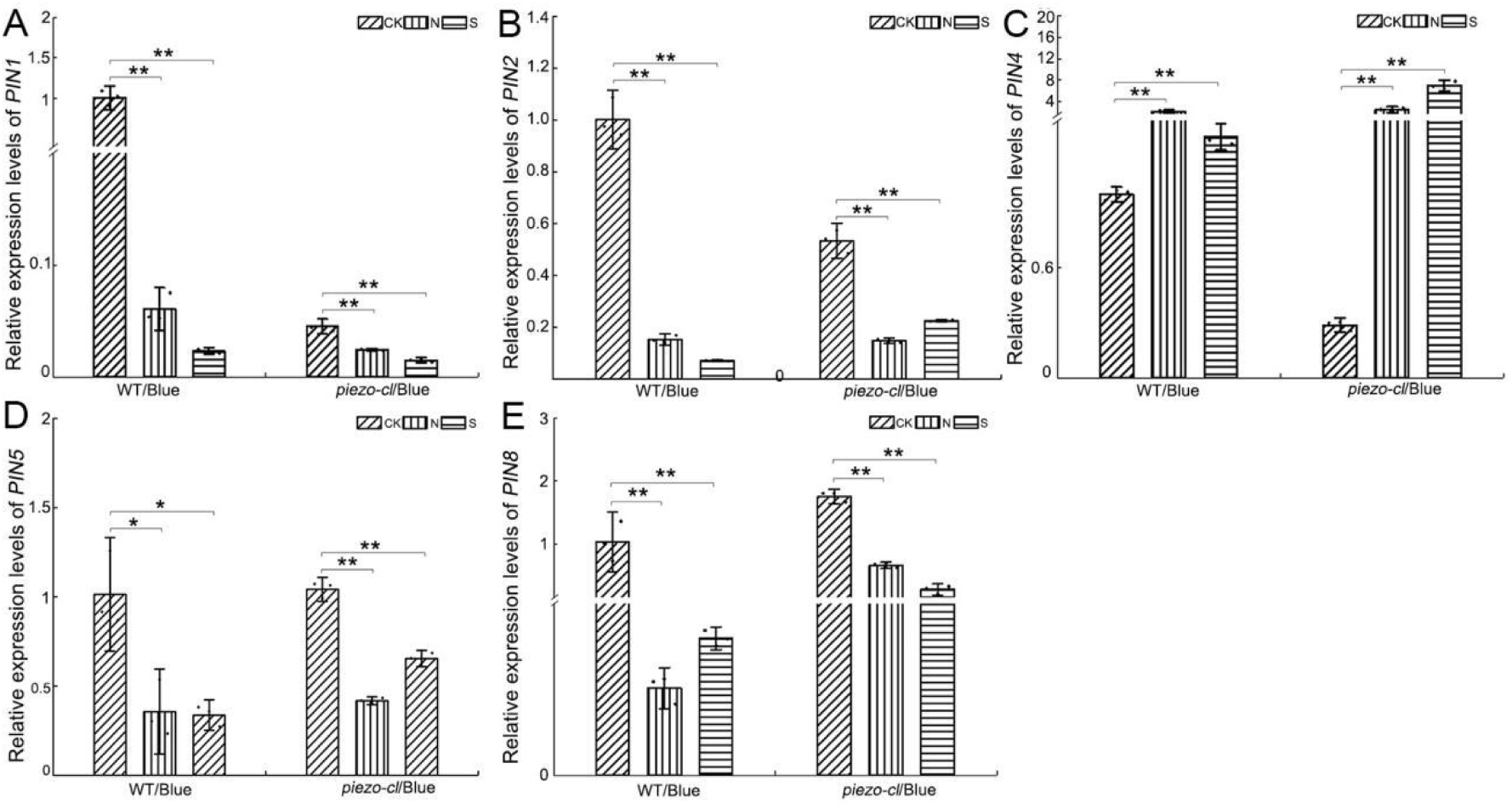
The expression levels of *PIN* genes in *piezo-cl* mutant seedlings subjected to a MF under blue light. The expression levels of *PIN1* (A), *PIN2* (B), *PIN4* (C), *PIN5* (D) and *PIN8* (E) genes in the roots of seedlings of WT and *piezo-cl* mutant lines subjected to a 500 mT MF under blue light. The *AtActin2* gene was used as an internal control. WT = wild-type. CK = seedlings not subjected to a MF, N = seedlings treated with the N pole of the magnetic block, S = seedlings treated with the S pole of the magnetic block. Blue = Seedlings treated with blue light. Data are means ± SD. The data presented here represent at least three biological replicates. ** *P* < 0.01 (Student’s *t*-test).

**Fig. S10.**
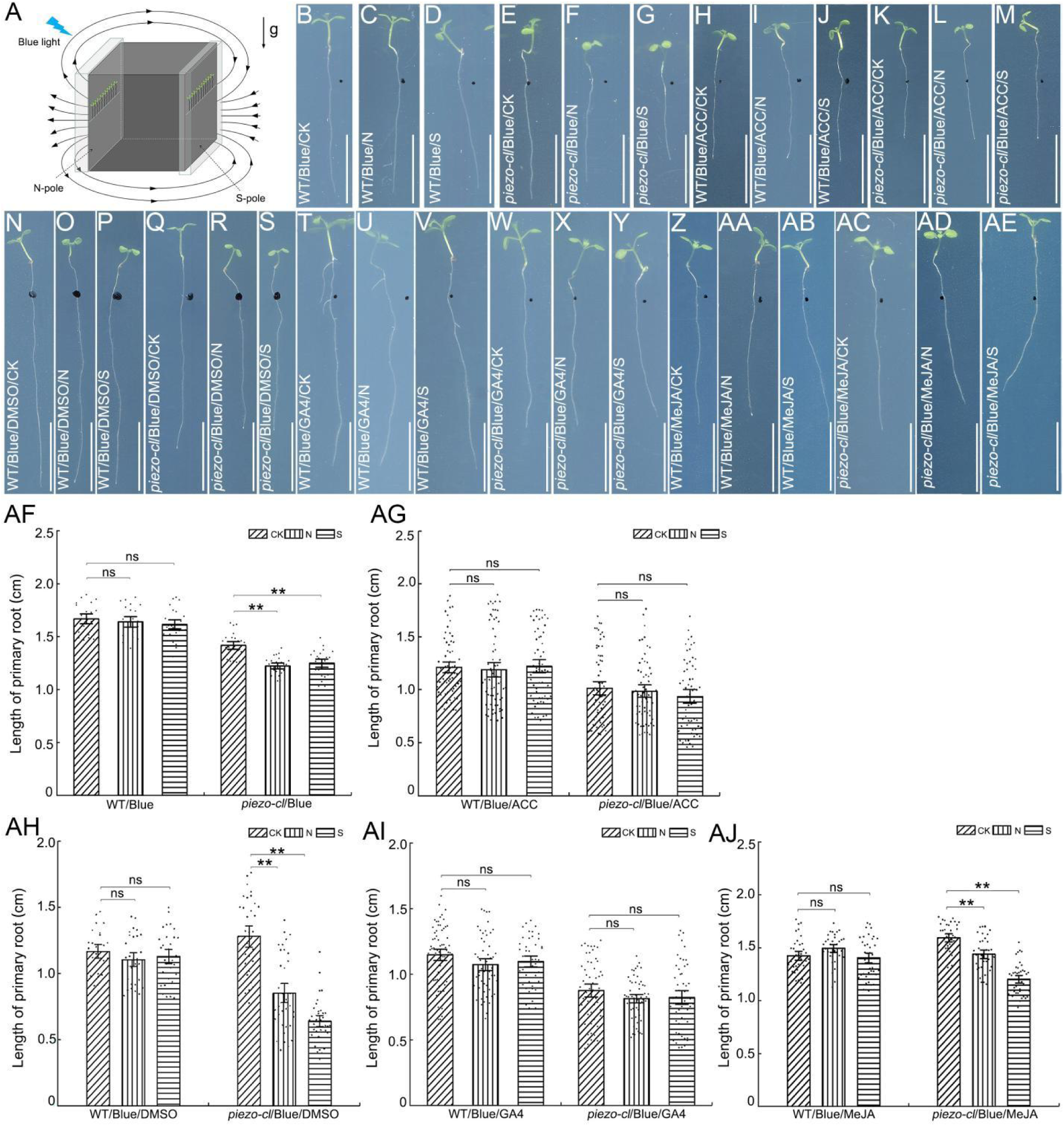
Root phenotypes of *piezo-cl* mutant seedlings treated with ACC, GA4, MeJA and MF under blue light. Schematic diagram of seedlings subjected to a MF under blue light (A). The root phenotypes of WT (B-D, H-J, N-P, T-V, Z-AB) and *piezo-cl* mutant (E-G, K-M, Q-S, W-Y, AC-AE) seedlings grown on 1/2 MS medium (B-G) as a control, or on 1/2 MS medium supplemented with 0.1 μM ACC (H-M), or 1/2 MS medium supplemented with DMSO (N-S) as a control, or with 0.1 μM GA4 (T-Y), or 0.1 μM MeJA (Z-AE) without a MF (B, E, H, K, N, Q, T, W, Z, AC) or subjected to a 500 mT MF (C-D, F-G, I-J, L-M, O-P, R-S, U-V, X-Y, AA-AB, AD-AE) under blue light (B-M, N-AE). Quantification of the primary root length of WT and *piezo-cl* mutant seedlings treated with ACC (AF-AG) (WT_1/2 MS_: n_CK_ = 21, n_N_ = 21, n_S_ = 25; *piezo-cl*_1/2 MS_: n_CK_ = 23, n_N_ = 23, n_S_ = 23; WT_ACC_: n_CK_ = 67, n_N_ = 65, n_S_ = 61; *piezo-cl*_ACC_: n_CK_ = 66, n_N_ = 61, n_S_ = 61), GA4 and MeJA (AH-AJ) (WT_DMSO_: n_CK_ = 23, n_N_ = 30, n_S_ = 23; *piezo-cl*_DMSO_: n_CK_ = 41, n_N_ = 42, n_S_ = 33; WT_GA4_: n_CK_ = 23, n_N_ = 30, n_S_ = 23; *piezo-cl*_GA4_: n_CK_ = 41, n_N_ = 42, n_S_ = 33; WT_MeJA_: n_CK_ = 23, n_N_ = 30, n_S_ = 23; *piezo-cl*_MeJA_: n_CK_ = 41, n_N_ = 42, n_S_ = 33). Arrows labeled “g” indicate the direction of gravity; the black, curved and solid lines with arrow heads represent the direction of the MF lines. The black dotted lines point to the N or S pole of the magnetic block, respectively. WT = wild-type. CK = seedlings not subjected to a MF, N = seedlings treated with the N pole of the magnetic block, S = seedlings treated with the S pole of the magnetic block. Blue = Seedlings treated with blue light. Data are means ±SE. ns = not significant, ** *P* < 0.01 (Student’s *t*-test). Scale bar = 1 cm.

**Fig. S11.**
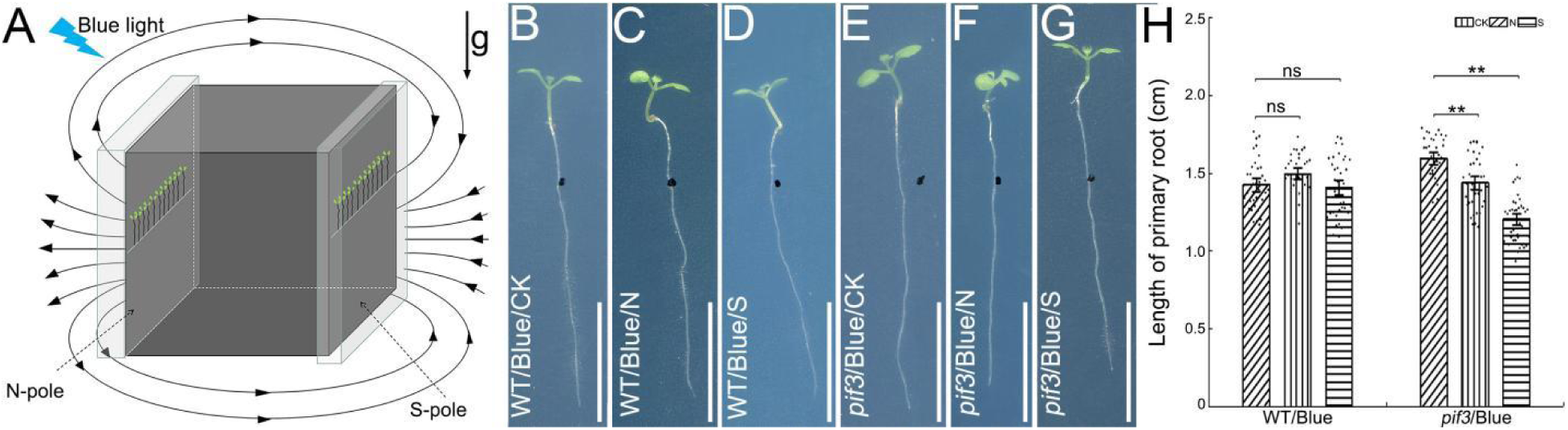
Root phenotypes of *pif3* mutant seedlings subjected to a MF under blue light. Schematic diagram of seedlings subjected to a MF under blue light (A). The root phenotypes of WT (B-D) and *pif3* mutant (E-G) seedlings not subjected to a MF (B, E), or subjected to a 500 mT MF (C-D, F-G) under blue light (B-G). Quantification of the primary root length of WT and *pif3* mutant (H) seedlings (WT: n_CK_ = 35, n_N_ = 33, n_S_ = 38; *pif3*: n_CK_ = 35, n_N_ = 37, n_S_ = 40). Arrows labeled “g” indicates the direction of gravity; the black, curved and solid lines with arrow heads represent the direction of the MF lines. The black dotted lines point to the N or S pole of the magnetic block, respectively. WT = wild-type. CK = seedlings not subjected to a MF, N = seedlings treated with the N pole of the magnetic block, S = seedlings treated with the S pole of the magnetic block. Blue = Seedlings treated with blue light. Data are means ± SE. ns = not significant, ** *P* < 0.01 (Student’s *t*-test). Scale bar = 1 cm.

**Fig.S12.**
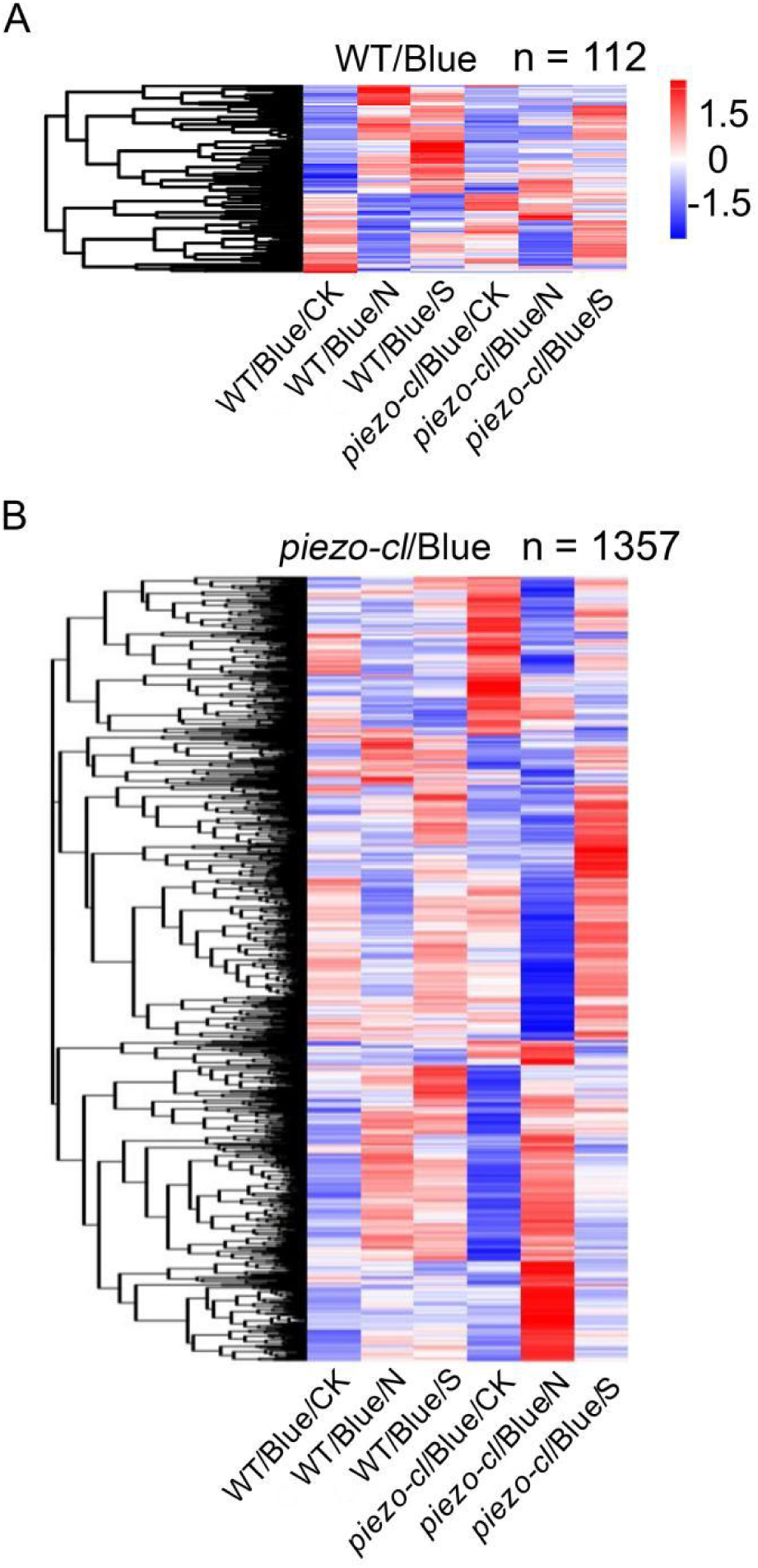
Heatmap of *Arabidopsis* unigenes in WT and *piezo-cl* mutant seedlings subjected to a MF under blue light. (A) Heatmap of DEGs in WT subjected to a MF under blue light, showing genes that exhibit either the same expression trend in response to both the N and S poles of the magnetic block or display specific changes in response to either the N or S pole of the magnetic block, compared to the control without MF treatment. (B) Heatmap of DEGs in *piezo-cl* seedlings subjected to a MF under blue light, showing genes that exhibit either the same expression trend in response to both the N and S poles of the magnetic block or display specific changes in response to either the N or S pole of the magnetic block, compared to the control without MF treatment. Gene expression levels of RNA-seq are log10 transformed (normalized FPKM + 1). “n” represents the number of DEGs. WT = wild-type. CK = seedlings not subjected to a MF, N = seedlings treated with the N pole of the magnetic block, S = seedlings treated with the S pole of the magnetic block. Blue = Seedlings treated with blue light.

**Fig.S13.**
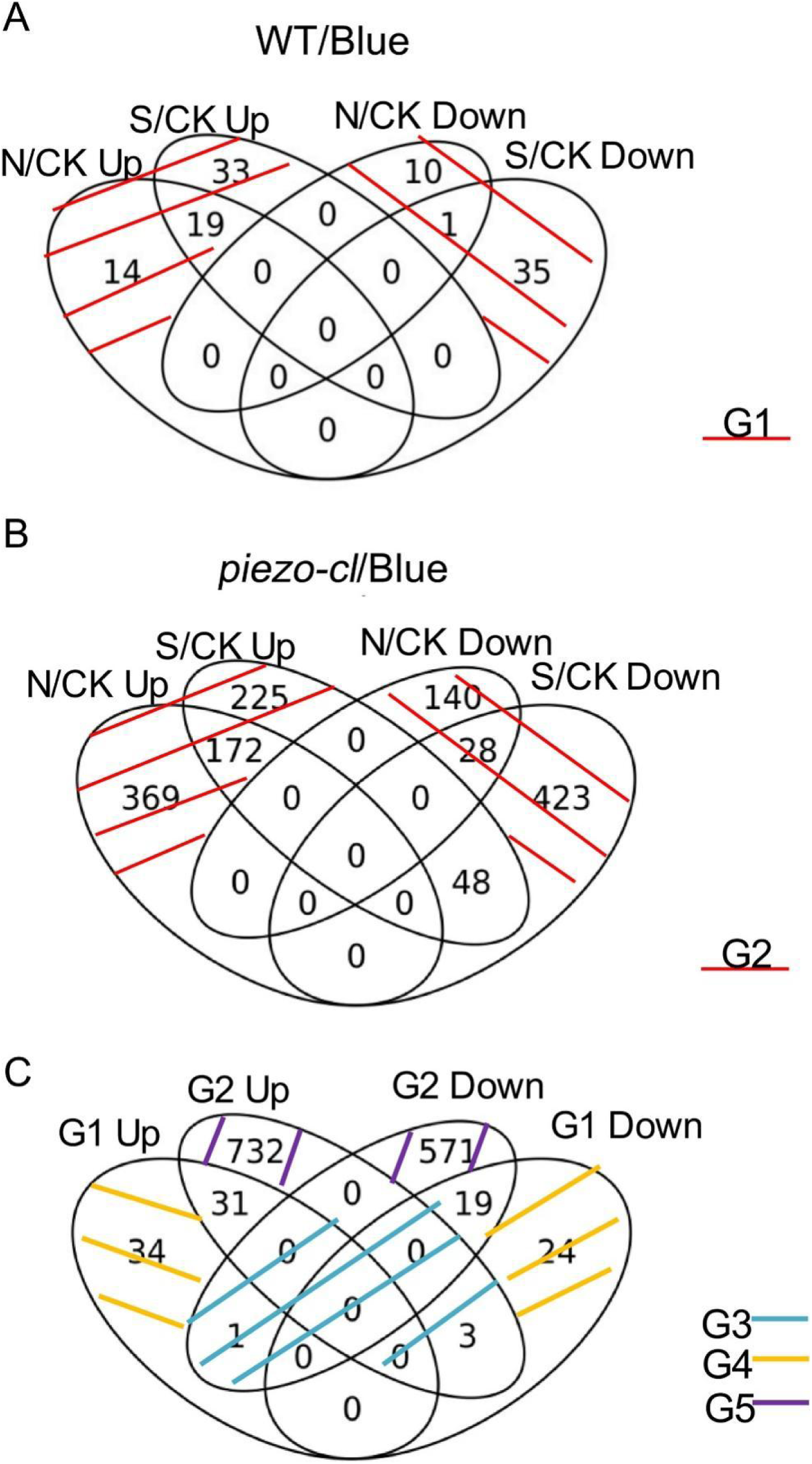
Expression patterns of DEGs in *Arabidopsis* seedling root transcriptomes. DEGs in WT seedlings exhibit similar trends under both N and S poles of the magnetic block, as well as show specific changes under the N or S pole of the magnetic block under blue light (G1) (A). DEGs in *piezo-cl* seedlings show consistent trends under both N and S poles of the magnetic block, with significant alterations specifically in response to either the N or S pole of the magnetic block under blue light (G2) (B). DEGs in seedlings grown in blue light that respond to MF regulation of root growth in both WT and *piezo-cl* mutant seedlings, but that exhibit different trends (G3), DEGs specifically responsive to MF treatment in WT seedlings (G4), and DEGs specifically responsive to MF treatment in *piezo-cl* mutant seedlings (G5) (C). WT = wild-type. CK = seedlings not subjected to a MF, N = seedlings treated with the N pole of the magnetic block, S = seedlings treated with the S pole of the magnetic block. Blue = Seedlings treated with blue light. up = up-regulated DEGs, down = down-regulated DEGs.

**Fig.S14.**
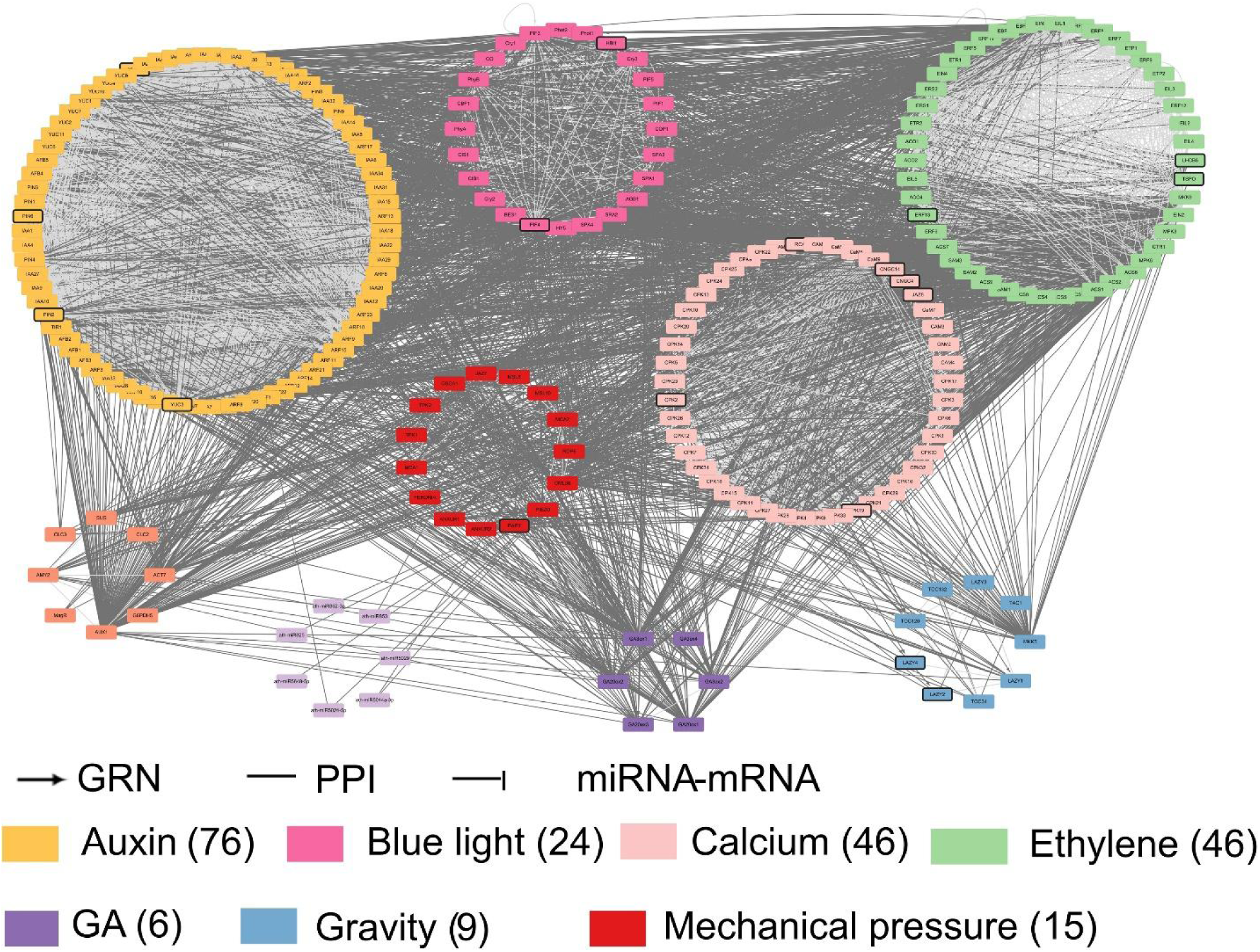
Integrated gene regulatory network (iGRN) in the roots of WT and *piezo-cl* mutant seedlings subjected to MF under blue light. The integrated network types include transcription factor regulatory networks (GRN), protein interaction networks (PPI) and miRNA-targets regulatory networks (miRNA-mRNA). The numbers in parentheses represent the counts of different types of genes.

**Table S2.** Phytohormone- and light-related responsive elements in the *PIEZO* promoter

| Name | Site | Strand | Organism | Sequences (5'-3') | Function |
| --- | --- | --- | --- | --- | --- |
| AE-box (47) | 1554 | + | *Arabidopsis thaliana* | AGAAACTT | Part of a module for light response |
| G-box (48-49) | 515 | - | *Arabidopsis thaliana* | TACGTG | Cis-acting regulatory element involved in light responsiveness |
|  | 1648 | + | *Zea mays* | CACGAC |  |
| P-box (50) | 1907 | + | *Oryza sativa* | CCTTTTG | Gibberellin-responsive element |
| TATC-box (51) | 1966 | - | *Oryza sativa* | TATCCCA | Cis-acting element involved in gibberellin-responsiveness |
| AuxRR-core (52) | 992 | - | *Nicotiana tabacum* | GGTCCAT | Cis-acting regulatory element involved in auxin |
| I-box (53) | 1968 | + | *Zea mays* | GGATAAGGTG | Part of a light responsive element |
| TCT-motif (54) | 645 | - | *Arabidopsis thaliana* | TCTTAC | Part of a light responsive element |
| GT1-motif (55) | 1802 | - | *Solanum tuberosum* | GTGTGTGAA | Light responsive element |
| GA-motif (56) | 321 | - | *Arabidopsis thaliana* | ATAGATAA | Part of a light responsive element |
| TCCC-motif (57) | 1123 | - | *Spinacia oleracea* | TCTCCCT | Part of a light responsive element |
|  | 1374 | - |  |  |  |
| GATA-motif (58) | 913 | + | *Solanum tuberosum* | AAGGATAAGG | Part of a light responsive element |
| CGTCA-motif (59) | 1415 | + | *Hordeum vulgare* | CGTCA | Cis-acting regulatory element involved in the MeJA-responsiveness |
| GARE-motif (60) | 298 | - | *Brassica oleracea* | TCTGTTG | Gibberellin-responsive element |
|  | 1299 | - |  |  |  |

**Table S3.**
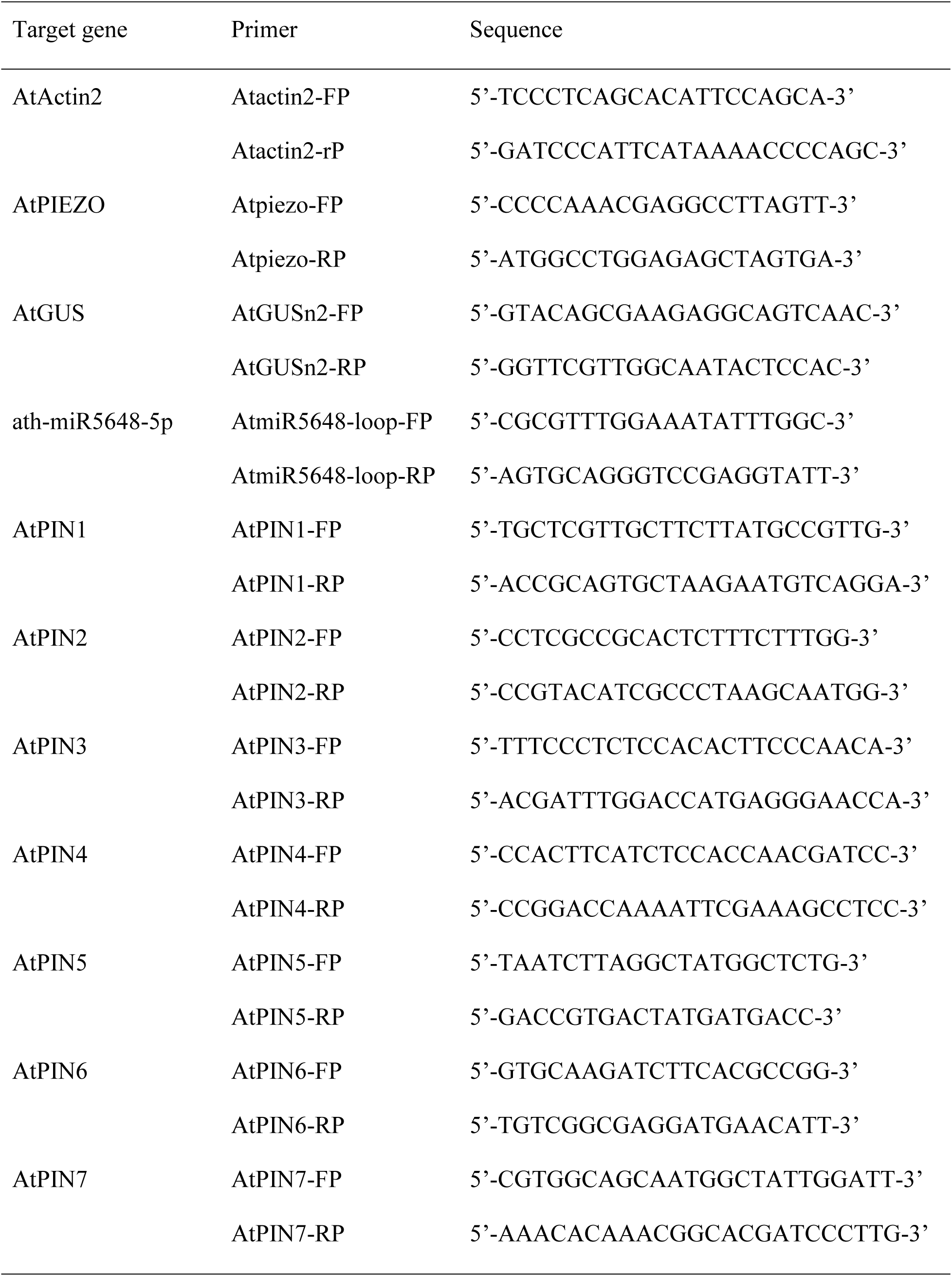
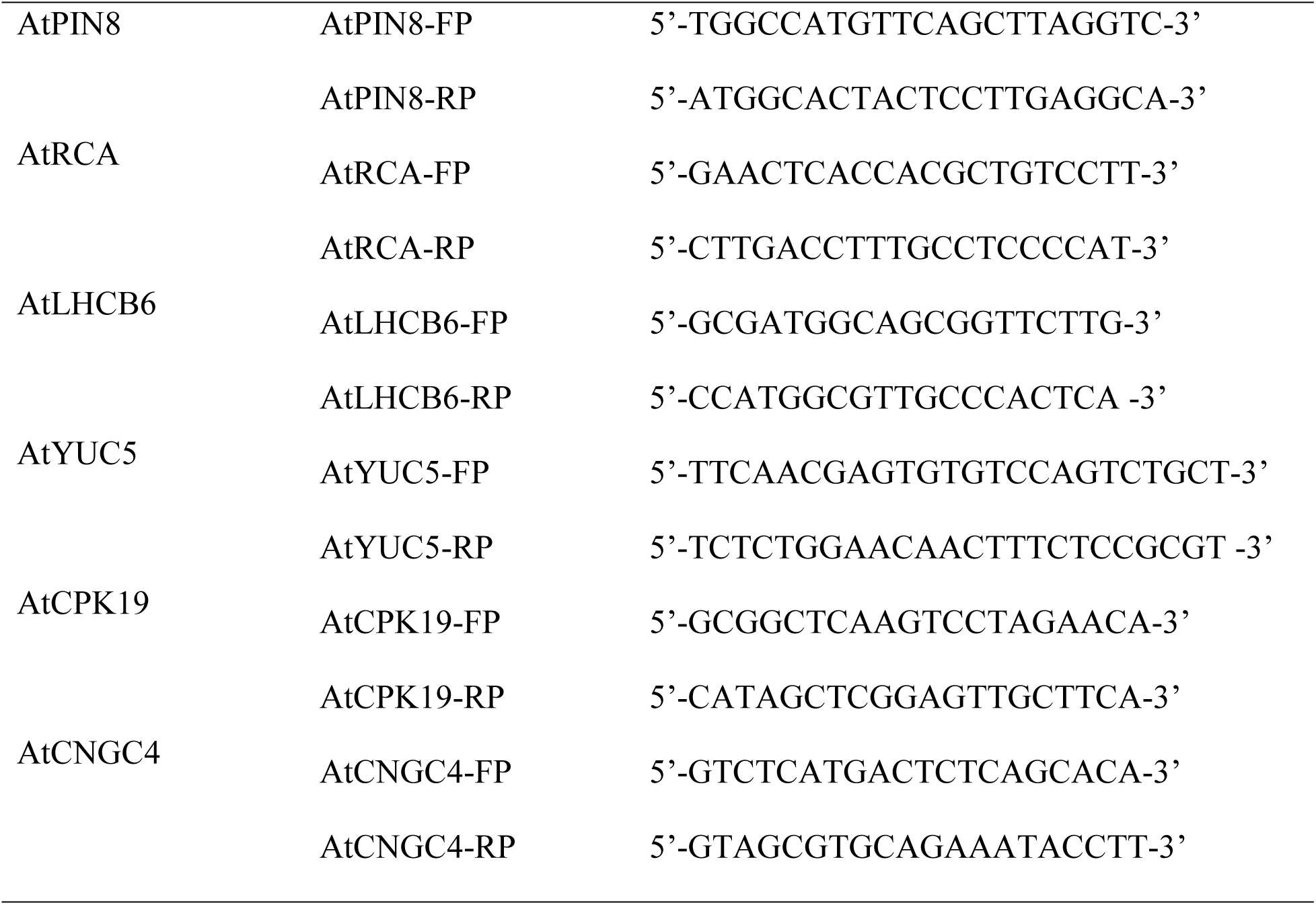
Sequences of gene-specific primers for real-time PCR

## Materials and Methods

### Plant materials and growth condition

The *Arabidopsis* lines used in this experiment were from an ecotype Columbia (WT) background, and included *piezo-c1* and *ProPIEZO::GUS* (*33*), *35S::PIEZO* lines, *piezo-T*, *cry_1-104_*, *cry_2-1_*, and *cry1/2* lines. The Arabidopsis *piezo-T* mutant line (NASCode:N663516) was purchased from AraShare (www.arashare.cn). *Arabidopsis* seeds were grown on 1/2 Murashige and Skoog (MS) medium (M5519; Sigma-Aldrich) containing 0.8 % agar and cold treated for 2 days in the dark at 4°C, and were then transferred to white light to grow 4 days at 22°C under 16 h light/8 h dark regime.

Four-day-old seedlings, where the primary root length was about 1 cm, were transferred to fresh 1/2 MS medium and tightly attached to the N and S poles on the surface of a NdFeB magnet N52 of strength 50, 200 or 500 mT (Hangzhou Permanent Magnet Group Co. Ltd, Hangzhou, China), in which the direction of the magnetic field (MF) was perpendicular to the direction of gravity. Seedlings were subjected to MF treatment under either red or blue light for 6 days. The blue light intensity was 96 μmol.m^-2^.s^-1^ and red light intensity was 48 μmol.m^-2^.s^-1^. Primary root lengths were measured using the ImageJ 1.41 software.

To detect the effect of the direction of the MF on regulating root growth, seedlings in which the primary root length with about 1 cm were transferred to a new plate with 1/2 MS medium, and then the plate was placed on either the N pole or the S pole of a NdFeB magnet N52 of strength 500 mT, in which the direction of the MF was either antiparallel or parallel to the direction of gravity. Seedlings were grown under MF treatment for 6 days under red or blue light.

To check the effect of a MF on the medium, the plate containing 1/2 MS medium was attached to the N or S pole of a NdFeB magnet N52 of strength 500 mT for 6 days in the dark. Four-day-old *Arabidopsis* seedlings grown on 1/2 MS medium under white light, and in which the primary root length was about 1 cm, were transferred to the 1/2 MS medium pre-treated with a MF at strength 500 mT, and grown under blue light for 6 days.

### *Arabidopsis* seedling root growth following treatment with auxin, GA, ethylene, MeJA and calcium ion inhibitor

Four-day-old seedlings in which the primary root length was about 1 cm were transferred to a plate containing 1/2 MS medium complemented with 5 μM of the auxin transport inhibitor TIBA, 0.01 μM NAA, 0.1 μM GA4, 100 nM ACC, 0.1 μM MeJA or 0.01 μM of the calcium ion inhibitor EGTA. The ACC and EGTA were dissolved in sterilized water. The chemicals TIBA, NAA, GA4 and MeJA were dissolved in DMSO, and seedlings treated with an equivalent volume of DMSO as a control. The plate containing the seedlings were attached to either the N or the S pole of a NdFeB magnet N52 of strength 500 mT and grown under blue light for 6 days.

### Root growth of seedlings grown with the roots in darkness

To observe the effect of blue light on the expression of the *PIEZO* gene, transgenic seeds with *ProPIEZO::GUS* were grown in a plate containing 1/2 MS medium. A black baffle was placed below the seeds at a distance of 1cm, and with a tiny space between the surface of the medium and the baffle to ensure that the seedling roots were able to grow through the baffle. A piece of tinfoil was used to cover the plate above the baffle to produce a dark environment, where the roots did not receive any light. Additionally, a piece of tinfoil was also used to cover the area that on the top of the seeds, so that light was prevented from directly irradiating into the space between the baffle and the medium, allowing the leaves to respond to light but keeping the roots in the dark. The seeds were sown on a plate with a baffle, tinfoil and the 1/2 MS medium, and were treated for 2 days at 4°C in darkness, and then the plates were transferred to blue light conditions and seedlings were grown for 4 days at 22°C. After 4 days, the plates together with the seedlings were transferred to a MF of strength 500 mT and grown for 6 days under blue light at 22°C.

### GUS staining

Ten-day-old seedlings were collected and stained with GUS solution [50 mM Na_2_HPO_4_·12H_2_O, 50 mM Na_2_HPO_4_·2H_2_O, 10 % (v/v) Triton X-100, 2 mM K_3_[Fe(CN_6_)], 2m M K_4_[Fe(CN_6_)]·3H_2_0, 10 mM EDTA, 1.04 mg/mL X-Gluc (R0851, Thermo Fisher Scientific)] at 37°C in darkness for 12 h. Then, the collected seedlings were then decolorized using ethanol. Samples were treated with ethanol at 30 %, 75 %, 90 %, and again 75 % (v/v) over 1 hour. Then, either the whole plant or just the root tip was observed and photographed using a Leica fluorescence microscope (DM2000, Leica, Germany).

### Calcium ion flux measurement

Root caps of 6-day-old seedlings of wild-type and *piezo-c1* mutant lines were used to measure Ca^2+^ flux using Non-invasive Micro-test Technology (NMT). Firstly, 100 mM CaCl_2_ filling solution was injected into the ion flow rate sensor (XY-CGQ-01) with a 4-5 μm aperture about 1.5cm long. Next, the Ca^2+^-LIX exchanger (XY-SJ-Ca-10) was taken up by the tip of the LIX Holder (XY-LIX-01), and then the tip of Ca^2+^-LIX exchanger was injected into the tip of the ion flow rate sensor with a syringe (XY-ZSQTZ-01) under the microscope. The Ca^2+^-LIX exchanger at the tip of the holder was injected into the tip of the ion flow rate sensor using a syringe (XY-ZSQTZ-01) under the microscope. Then, a silver wire was chlorinated in silver chloride (XY-RY-05) for 25 seconds until it was an off-white color, and then inserted into the ion flux rate sensor, and the mounted ion flow rate sensor was connected to the pre-ion amplifier. The prepared Ca^2+^ flow rate sensors were also calibrated by the three-point calibration method using calibration solution 1, calibration solution 2 (XY-RY-02), and 0.1 mM CaCl_2_ test solution (XY-RY-01) with a theoretical value of 27 ± 5 mv/decade. The roots of seedling wild-type and *piezo-c1* lines were fixed to the bottom of a 35 mm petri dish (XY-PYM-35) using resin block (XY-SZK). 0.1 mM CaCl_2_ (XY-RY-01) was added to submerge and stabilize the samples for 15 min, and then the samples were placed on the NMT microscope stage, the reference electrode (YG-CBDJ-01) was inserted into a 0.1 mM CaCl_2_ solution (XY-RY-01), and the stage and preion amplifier were adjusted to position the Ca^2+^ flux rate sensor near the root cap of the primary root. Finally, the N or S pole of a magnetic rod of strength 120 mT was placed close to and above the root cap, and the blue filters of the NMT microscope were used as the blue light source. After root samples were stabilized in the test solution for 20 min, the Ca^2+^ flux was began to assay in a time course of 10 min for each data collection. Flow rate data were collected directly using the imFluxes V3.0 software (Xuyue.net) in picomole•cm^-2^-s^-1^, with positive values indicating Ca^2+^ efflux and negative values indicating Ca^2+^ influx. At least 7 roots were tested in each experiment for both the wild-type and *piezo-cl* mutant seedlings. All the reagents and consumable materials used in the experiments were purchased from Beijing Xuyue Technology Co. Ltd (Beijing, China).

### RNA isolation and cDNA synthesis

The primary roots of 6-day-old wild-type or *piezo-c1* mutant seedlings that had been subjected to a 500 mT MF for 15 min, 30 min, 72 h or 144 h under blue light were collected. The primary roots of 6-day-old wild-type seedlings or seedlings expressing *ProPIEZO::GUS* subjected to 500 mT MF treatment for 6 days were used to isolate RNA. The isolation of RNA from the primary roots and cDNA biosynthesis referred to a previous report (*43*).

### Real-time PCR analysis

The relative quantitative gene expression levels were determined using an ABI QuantStudio 7 Flex Real-Time PCR System (Applied Biosystems, USA). The 10 μL PCR reaction mixture contained 5 μL NovoStart ^Ⓡ^ SYBR qPCR SuperMix Plus (Novoprotein Scientific lnc, Shanghai), 1 μL of cDNA template, 0.2 μL ROX2 (Novoprotein Scientific lnc, Shanghai), 3 μL RNase free H_2_O and 0.8 μL of a 10 μM solution containing one of the following primer pairs: Atpiezo-FP and Atpiezo-RP, AtGUSn2-FP and AtGUSn2-RP, AtmiR5648-loop-FP and AtmiR5648-loop-RP, AtPIN1-FP and AtPIN1-RP, AtPIN2-FP and AtPIN2-RP, AtPIN3-FP and AtPIN3-RP, AtPIN4-FP and AtPIN4-RP, AtPIN5-FP and AtPIN5-RP, AtPIN6-FP and AtPIN6-RP, AtPIN7-FP and AtPIN7-RP, AtPIN8-FP and AtPIN8-RP, AtRCA-FP and AtRCA-RP, AtLHCB6-FP and AtLHCB6-RP, AtCPK19-FP and AtCPK19-RP, AtYUC5-FP and AtYUC5-RP, or AtCGNC4-FP and AtCGNC4-RP, which were used to amplify the genes *AtPIEZO, GUS, ath-miR5648-5p*, *AtPIN1, AtPIN2, AtPIN3, AtPIN4, AtPIN5, AtPIN6, AtPIN7, AtPIN8, AtRCA*, *AtLHCB6*, *AtCPK19*, *AtYUC5* and *AtCGNC4*, respectively (Supplemental Table S3). The *AtActin2* gene (AT3G18780) acted as the internal control, and was amplified with the primer pair AtActin2-FP and AtActin2-RP (Supplemental Table S3). PCR was performed under the following conditions: denaturation at 95°C for 2 minutes, followed by 40 cycles of 95°C for 45 seconds, 55-62°C for 30 seconds and 72°C for 1 minute. Three biological replicates were made. The relative gene expression levels were calculated by using the 2^−ΔΔCt^ method. The software SPSS version 19.0 (IBM, Inc., Armonk, NY, USA) was used to analyze differences in gene expression. * *P* < 0.05 was taken to indicate a statistical difference, ** *P* < 0.01 indicated a statistically significant difference.

### Transcriptome analysis

Both of the raw RNA-seq and sRNA-seq reads were subjected to quality control with FastQC (v0.12.1). Subsequently, transcriptome analysis was performed using the HISAT2 pipeline (*44*). The *Arabidopsis* genome version used was TAIR10, downloaded from the phytozome v13 (https://phytozome-next.jgi.doe.gov/). The differentially expressed genes were identified using GFOLD (*45*)

### Construction of the integrated gene regulatory network

The integrated gene regulatory network encompasses three types of networks: the transcription factor regulatory network, with a score of more than 0.7 (Nature Plants 2021), the protein-protein interaction network (https://string-db.org/), with a score of more than 0.7 and the miRNA-target gene regulatory network. The regulation of target genes by miRNAs was predicted with psRNATarget (Nucleic Acids Research 2018). Sub-network was extracted with Ctyohubba tool and visualized in Cytoscape (v3.9.0).

### Promoter analysis of *AtPIEZO* gene

The website Ensembl Plants website (http://plants.ensembl.org/index.html) was searched for the *AtPIEZO* (AT2G48060) sequence, and the 2000 bp upstream of the *AtPIEZO* gene was used to analyze the promoter structure using online software Plant Care (http://bioinformatics.psb.ugent. be/webtools/plantcare/html/). A visual representation was constructed using TBtools (*46*).

### Statistical analysis

The ImageJ 1.41 software was used to measure root length, root growth angle and GUS level. All images were processed using the Photoshop software. Data were analyzed with one-way ANOVA or two-tailed Student’s *t*-test, and the Origin 2022 and Photoshop were used to plot images.

